# Mouse olfactory system acts as anemo-detector and -discriminator

**DOI:** 10.1101/2024.08.28.610087

**Authors:** Sarang Mahajan, Suhel Tamboli, Susobhan Das, Anindya S. Bhattacharjee, Meenakshi Pardasani, Priyadharshini Srikanth, Shruti D. Marathe, Avi Adlakha, Lavanya Ranjan, Sanyukta Pandey, Nixon M. Abraham

## Abstract

Airflow detection while smelling is a fundamental requirement for olfaction, yet the mechanisms underlying such multimodal processing in the olfactory system remain unknown. We report here that mice can learn to accurately discriminate airflow with parallel processing of both mechanical and chemical stimuli revealed by modulated sniffing and refined calcium signaling in the olfactory bulb inhibitory network. Genetic perturbation of AMPAR function and optogenetic control bidirectionally shifted airflow discrimination learning pace, with contrasting phenotypes observed for odor learning, engagement of inhibitory circuits, and setting the optimal inhibition level for stimulus refinement. Multimodal odor-airflow stimuli at subthreshold levels enhanced learning, demonstrating that mechanical stimuli heighten olfactory perception. Our results, thus explain the multimodality of olfaction, and reveal an unexplored dimensionality of odor perception.

## Introduction

The multi-sensory environment renders the brain to perceive the external world by combining concurrently occurring diverse sensory stimuli. While humans rely vastly on visual and auditory inputs, rodents primarily use olfactory and vibrissal cues from their surroundings. In nature, odor plumes carry odorant molecules, which are associated with varying airflows. What is the role of carrier airflow in olfactory perception? The hypothesis of mechanosensation through the olfactory system emerged from the seminal works of Adrian and Domino (Adrian, 1951; Ueki and Domino, 1961). Further evidence from the olfactory bulb (OB) recordings, reporting the neural activity even in the absence of odorant stimuli, strengthened this hypothesis (Macrides and Chorover, 1972; Onoda and Mori, 1980a). Recently, this hypothesis was tested by recording electrical activity from OSNs in the presence of mechanical stimuli (Connelly et al., 2015; Grosmaitre et al., 2007). OSNs in the rodent nose detect and transfer mechanical (Connelly et al., 2015; Grosmaitre et al., 2007) and odorant information to the OB (Buck and Axel, 1991; Mombaerts et al., 1996). In the OB, the signaling through mitral and/or tufted cells (MTCs) is refined by an inhibitory network (Abraham et al., 2010; Gire and Schoppa, 2009) and top-down modulatory inputs (Fletcher and Chen, 2010). Hence, OB may act as the first site of integration of odorant and airflow-related information in mammals. Despite existing literature on cellular mechanisms of inhibitory interactions (Abraham et al., 2010; Gire and Schoppa, 2009; Gschwend et al., 2015), their functional relevance in the context of combined olfactory and mechanosensory information processing remains largely unknown.

Sniffing is an integral part of olfactory perception, not only to gather information of volatiles from the surroundings of humans and animals, but even to communicate socially relevant information such as the hierarchical status in the animal kingdom (Abraham et al., 2014; Mainland and Sobel, 2006; Uchida et al., 2006; Wachowiak, 2011; Wesson, 2013). The respiration-coupled activities in different centers of the olfactory pathway have been reported from the time of Adrian’s seminal experiments (Adrian, 1942). While the odor representation observed in the OB (Bathellier et al., 2008a; Cury and Uchida, 2010; Gschwend et al., 2012; Smear et al., 2011), and the piriform cortex (Miura et al., 2012) can be sniffing-dependent, what information could be coded by the modulations of respiration in the absence of odorants? Olfactory perception is tightly coupled to the respiratory pattern (Cury and Uchida, 2010). The rhythmic activity of the neurons in the Pre-Bötzinger complex may control the mechanical drive needed for active sampling of odorants (Kleinfeld et al., 2014; Smith et al., 1991). Indeed, the oscillatory activity in the OB projection neurons can be generated by the sniffing-induced mechanosensation, in a glomerulus-specific manner. This has been shown to be critical for phase coding of odor identity information (Iwata et al., 2017). Despite these experimental evidence, the neural mechanisms underlying mechanosensation through OB circuits remain undiscovered.

How do OB circuits maintain a robust phase code? What is the role of OB inhibitory circuits in regulating the mechanosensation through olfactory system? The activity patterns elicited in the OB by airflow are broadly distributed compared to the odor-evoked activity patterns (Wu et al., 2017). This implies a larger activation of the inhibitory network in response to airflow signals. A specific odorant molecule activates a subset of OSNs, thereby resulting in specific odor-evoked glomerular activity patterns. OB inhibitory circuits modulate the spiking activity of projection neurons and help refine the odorant information that is conveyed to higher centers (Abraham et al., 2010; Gschwend et al., 2015). As airflow evokes broader activation, the optimized inhibition needed for refining the information would be different compared to odor information. Hence, probing the role of inhibitory network is pivotal for understanding the fundamental principles of mechanosensation through the rodent olfactory system.

Here, we show that mice can detect and discriminate ethologically relevant airflow rates through their nose. The in vivo Ca^2+^ imaging from inhibitory interneurons, and the modulation of inhibitory circuits using genetic (deletion of GluA2 subunit of AMPA receptors) and optogenetic approaches (ChR2 and Arch activations) confirmed the role of OB circuits in processing and refining airflow information. We also elucidated that the optimal inhibition needed for refining odor vs. airflow discrimination differed significantly, indicating the varying extent of inhibitory circuit recruitment by chemical and mechanical stimuli. Furthermore, we show that mechanical information processing through OB circuits can facilitate olfactory perception at subthreshold levels. Together, these results prove that the mouse olfactory system can act as anemo-detector and -discriminator.

## Results

### Mice can detect and discriminate diverse airflow rates using their nose, even in the absence of whiskers

The less-explored dimensions and the boundless chemical world make olfactory stimulus space challenging to resolve (Bushdid et al., 2014; Gerkin and Castro, 2015; Meister, 2015). Given the probable variations of odor stimuli with the mechanical (airflow-induced) and thermal (for example, volatility changes by temperature) components in nature make the olfactory perception unique. To investigate how mechanosensation modulates olfactory perception, we started by studying mice’ discrimination behavior in the absence of any odor molecule. To achieve this, we built a customized instrument capable of providing airflows as stimuli in three different modes (M1, M2, and M3 modes, Figure 1). As airflow stimuli can be perceived through mice’ whisker system, these modes of stimulus delivery were decided to enable whisker activation. We started training the animals to discriminate the airflows using M1 mode, 8 liter per minute (LPM) vs. 4 LPM [airflow (4 s) is delivered from the top, Figure 1A]. These flowrates were powerful enough to cause deflection of a plucked whisker. Animals reached the asymptotic levels of learning (accuracy >85%) within 900 trials (Figure 1B1). Having observed their discrimination abilities, we continued the training on various airflow pairs to optimize the stimulus strength and duration. Mice successfully learned to discriminate different airflow pairs within 600-900 trials, reaching ∼90% accuracy towards the end of training (2 LPM vs. 1 LPM, 3 s duration; 0.8 LPM vs. 0.3 LPM, 3 s duration; 0.8 LPM vs. 0.3 LPM, 2 s duration, Figure 1B1, B2). To rule out the possibility of using any nonspecific cues while discriminating airflows, mice were trained on a control task (0.3 vs. 0.3 LPM, 2s duration), wherein they performed at the chance level (Figure 1B1, 1B2). This was further confirmed by the photoionization detector (PID) analysis of various airflows and odor stimuli. We did not observe any significant voltage shifts for airflows, whereas a well-defined voltage shift was observed during the odor stimulus (Figure 1B3, Supplementary Figure S1). Animals were optimally motivated to perform these tasks, which were quantified by inter-trial intervals (Figure 1C). As mice performed discrimination of different flow pairs with high accuracy, we quantified another behavioral readout, i.e. time taken by mice to discriminate various flow pairs. This was calculated from the lick patterns shown by animals towards rewarded and non-rewarded airflow stimuli (Figure 1D1). Our results show comparable flow discrimination time (FDT) across different airflow pairs, making the sensing mechanism and the underlying neural pathways an interesting open question (Figure 1D2, one-way repeated measures ANOVA, F = 0.4158, p = 0.6725).

**Figure 1.**
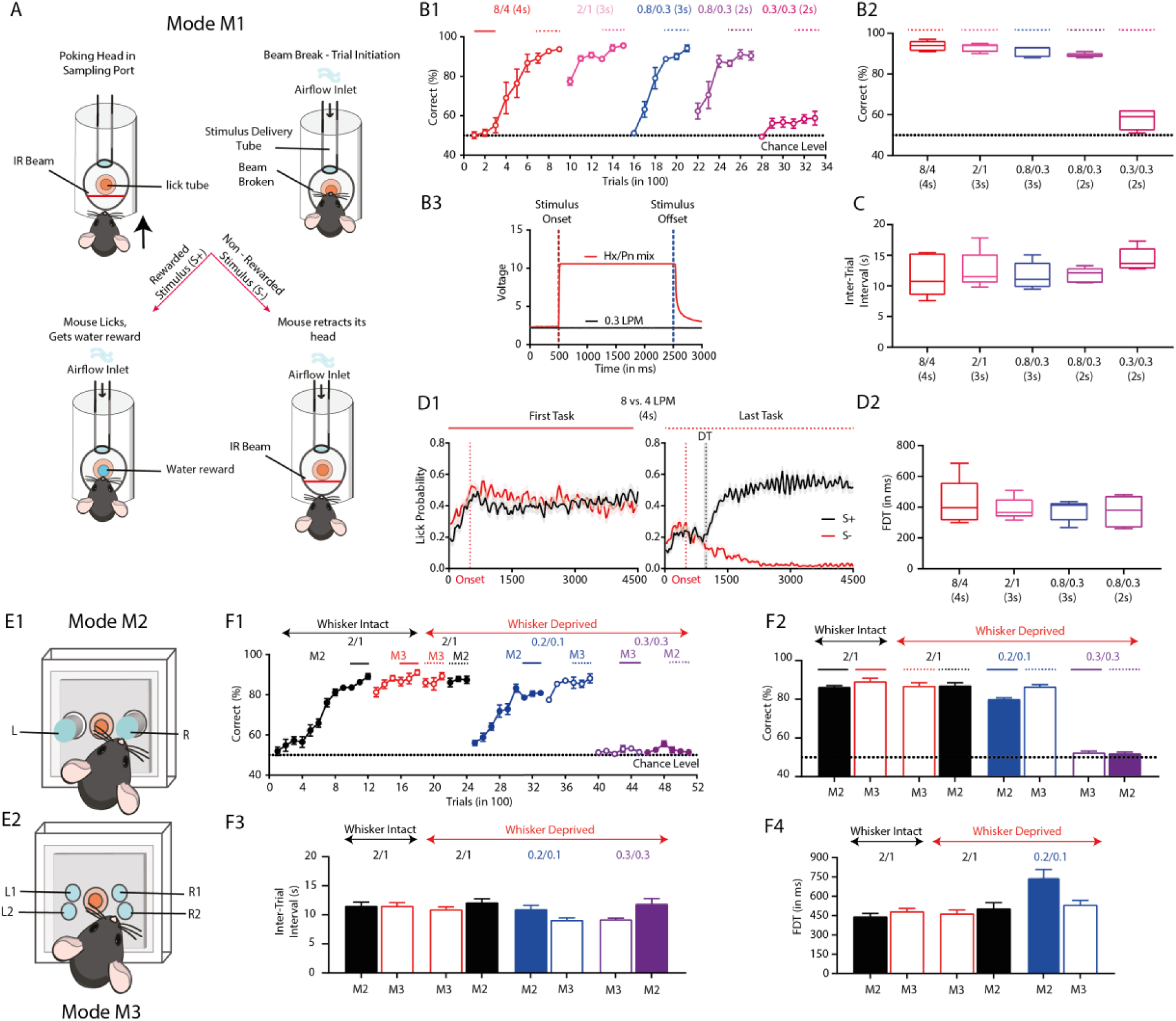
Mice can detect and discriminate airflows in the absence of whiskers. **A.** Go/No-Go operant conditioning paradigm for M1 Mode. A trial is initiated as soon as the mouse pokes its head into the sampling port guarded by the IR beam. Animal receives the stimulus and samples it. If the stimulus is rewarded (S+), animal licks on the lick tube and the correct licking response triggers the delivery of the water reward. If the stimulus is unrewarded (S–), a trained animal doesn’t lick and retracts its head from the sampling port. **B.** Animals can detect and discriminate different airflow pairs in 2s stimulus duration. **B1.** Airflow discrimination learning accuracies measured for diverse airflow pairs at different stimulus durations. Each point on the learning curve represents the average of 100 trials across all animals (n = 5). The black dotted line at y = 50 represents the chance level. **B2.** Learning accuracies measured from the last 300 trials for various airflow pairs at different stimulus durations and averaged across animals. The final accuracies were found to be greater than 80% for all different stimulus durations except for 0.3 vs. 0.3 LPM where accuracies were at around 50% (chance level). Final accuracies for 8 vs. 4 LPM (4s) - 93.80 ± 1.068 [Mean ± SEM]; 2 vs. 1 LPM (3s) - 93 ± 0.8944; 0.8 vs. 0.3 (3s) LPM - 91.2 ± 1.114; 0.8 vs 0.3 LPM (2s) - 89.2 ± 0.4899; 0.3 vs. 0.3 LPM (2s) - 57.60 ± 2.2205. B3. PID profile of airflow and odors reveal absence of any odor stimulus in the airflow. Comparison of PID profiles of airflow (0.3 LPM - black) and odor (Hexanal-pentanone mixture - red). After stimulus onset the airflow PID profile remains flat, however, a sharp and sustained increase in the voltage for odor duration was observed. **C.** Inter-trial intervals (ITIs) measured from the last 300 trials for different airflow pairs and stimulus durations. The average ITIs for all airflow pairs: 8 vs. 4 LPM (4s) - 11.67 ± 1.536 [Mean ± SEM]; 2 vs. 1 LPM (3s) - 12.57 ± 1.378; 0.8 vs. 0.3 (3s) LPM - 11.67 ± 0.9778; 0.8 vs 0.3 LPM (2s) - 11.78 ± 0.5339; 0.3 vs. 0.3 LPM (2s) - 14.33 ± 0.8282. The ITIs were in the range of 11-14 s which is the optimal range of the motivation of animals. **D.** FDT for airflow discrimination tasks are calculated from the lick patterns. **D1.** Representative lick responses of a single animal averaged over 300 trials (150 S+ and 150 S–) during first and last task i.e. initial and final phases of learning, respectively. Red traces represent average of S+ trials where as black traces represent average of S– trials. The shaded region represents SEM. The red dotted vertical line represents the onset of the stimulus. During the first task, as the animal did not learn the discrimination task and performed at chance level, the lick responses for S+ and S– trials were similar. During the last task, as the animal performed with higher accuracy, the lick responses for S+ and S– trials diverged. Statistical difference is calculated between 150 S+ and 150 S– trials and the last time point where the p < 0.05 is considered as FDT. The black dotted vertical line represents the average FDT and the shaded area represents the SEM. **D2.** Average FDTs for different airflow pairs were in the range of 300-500 ms and were similar for all the airflow pairs (8 vs. 4 LPM (4s) - 428.0 ± 67.88 ms [Mean ± SEM]; 2 vs. 1 LPM (3s) - 388.0 ± 32.20; 0.8 vs. 0.3 (3s) LPM - 378.4 ± 29.98; 0.8 vs 0.3 LPM (2s) - 372.0 ± 44.99; one-way repeated measures ANOVA, F = 0.4158, p = 0.6725, n=5 animals). **E.** Different modes of stimulus delivery. In Mode M2 **(E1)** and Mode M3 **(E2)** the stimulus is delivered on the whiskers of the animal in a focused and diffusive way, respectively. **F.** Animals can discriminate the airflow pairs in the absence of whiskers. **F1.** Training schedule and airflow discrimination accuracies measured for different airflows under different conditions. The conditions used were 2 vs. 1 LPM in M2 mode followed by M3 mode. The whiskers were then trimmed and animals’ performance was tested on 2 vs. 1 LPM for M3 mode followed by M2 mode. The animals were further trained on different airflow pairs including 0.2 vs. 0.1 LPM and 0.3 vs. 0.3 LPM for both M2 and M3 mode of stimulus delivery under whiskers-deprived conditions. The learning of animals remained independent of the stimulus delivery mode as well as the intactness of whiskers. **F2.** Learning accuracies measured from last 300 trials for different airflow pairs across different conditions, as shown in F1. The final accuracies for different airflows were found to be > 80% for both M2 and M3 mode under whiskers intact and deprived conditions except 0.3 vs. 0.3 LPM where learning accuracies were near the chance level. WI vs. WD comparison - M2 mode: two-tailed paired t-test, p = 0.6135, M3 mode: two-tailed paired t-test, p = 0.3024. **F3.** Inter-trial intervals (ITIs) measured from last 300 trials for different airflow pairs and different modes. The average ITIs was in the range of 9-12 s for different airflow pairs under different conditions. **F4.** Average FDTs for different airflow pairs under different stimulus delivery and whisker-deprived conditions were in the range of 300-500 ms. The FDTs before and after whisker deprivation for both modes were similar (M2 mode: whiskers intact (WI) - 440 ± 27.52 ms, whiskers deprived (WD) - 502.9 ± 46.89 ms, two-tailed paired t-test, p = 0.3589, M3 mode: WI - 480 ± 25.91 ms, WD - 462.9 ± 30.99 ms, two-tailed paired t-test, p = 0.6627).

Having observed mice’ airflow discrimination abilities, we tried to deliver the airflow stimulus dissociated from the reward delivery tube to monitor their sampling strategies with more precision. We delivered the flows from two different focal points in one mode (M2, Figure 1E1) and in a more diffused manner in the second mode (M3, Figure 1E2) and trained the animals to discriminate 2 vs. 1 LPM. Mice were able to reach the asymptotic phase of learning within 1200 trials and the final accuracies remained similar for both M2 and M3 mode (Figure 1F1, F2: M2 mode - 86.33 ± 0.7182, M3 mode - 89.21 ± 1.650, two-tailed paired t-test, p = 0.1348). From our video analysis we observed mice using their nose to sample the airflows during the airflow discrimination task. As we observed this specific sampling behavior, we blocked the possible airflow information processing through whiskers by trimming them and challenging the mice to discriminate 2 LPM vs. 1 LPM. Mice were able to discriminate these airflows with high accuracy in the absence of whiskers (Figure 1F1, F2: M2 mode: whiskers-intact (WI) - 86.33 ± 0.7182, whiskers-deprived (WD) - 87.14 ± 1.438, two-tailed paired t-test, p = 0.6135, M3 mode: WI - 89.21 ± 1.650, WD - 86.90 ± 1.628, two-tailed paired t-test, p = 0.3024). We further continued the training using airflows of lower strength, i.e. 0.2 LPM vs. 0.1 LPM, using different modes, where whisker-deprived mice reached higher accuracy in a few hundred trials. Mice failed to discriminate when they were challenged with the same airflow rates (0.3 vs. 0.3 LPM, Figure 1F1, F2). The efficacy of animals discriminating airflow rates in the absence of whiskers was also demonstrated by another method of whisker deprivation i.e. whisker plucking, wherein the performance of animals was not different before and after the deprivation (Supplementary Figure S2). These results confirm mice’ ability to discriminate various airflow rates in the absence of whiskers and using their nose (Figure 1F1). High accuracy performances were observed in WI and WD mice at the end of each training session except the chance level performance observed for the control task that involved the discrimination of same airflow rates (Figure 1F2). Mice showed optimal motivation levels, quantified by ITIs, under WI and WD conditions (Figure 1F3). We further analyzed the FDTs from various airflow discriminations and found similar FDTs in WI and WD mice, ruling out the possible involvement of whisker system in the observed anemo-discrimination behavior (Figure 1F4, M2 and M3 mode, before and after whisker deprivation, two-tailed paired t-test, p = 0.3589 and p = 0.6627, respectively).

### Olfactory information processing centers are involved in airflow rate discrimination

As we demonstrated the involvement of rodent nose in discriminating a large range of airflow rates, next we tried to delineate the role of olfactory centers in processing ethologically relevant airflow rate information. As a first step, we quantified the airflow distribution from rodent habitats in different locations using a hot wire anemometer. Our measurements from 40 different locations showed that 90% of measured airflow lay in the range of 0.1 - 0.6 LPM (Figure 2A). This airflow range was mostly considered further for designing the stimuli pairs in anemo-discrimination experiments. The role of olfactory epithelium in airflow rate discrimination was investigated by ablating olfactory sensory neurons (OSNs) using Zinc sulphate (ZnSO_4_) intranasal- and methimazole (MMZ, an olfactotoxic drug) IP-injections as both of these treatments have shown to induce hyposmia/anosmia in animals (Alberts and Galef, 1971; Bergman and Brittebo, 1999) A significant reduction in the thickness of OSN layer was observed two days after the treatments, indicating the efficiency of these methods in ablating OSNs (Figure 2B, 2C, Vehicle - 49.95 ± 1.110, Methimazole - 18.24 ± 1.035, ZnSO_4_ - 17.41 ± 0.7997, one-way ANOVA, F = 354.2, p < 0.0001). To study the involvement of OSNs in anemo- discrimination, three batches of mice were trained on two airflow discrimination tasks (0.2 vs. 0.1 LPM, and 0.6 vs. 0.3 LPM) and an odor discrimination task (Amyl acetate, AA vs. Ethyl butyrate, EB, using same flow rate, 0.6 LPM). All groups of mice learned to discriminate the airflow rates and odors with similar accuracy (Figure 2D1, two-way ANOVA with Tukey’s multiple comparison, p>0.05). These mice were then administered with the following: 5% ZnSO_4_ (intranasal application, Group 1), MMZ (IP injection, Group 2), and Phosphate buffer saline (PBS, intranasal application and IP injections, Group 3). The performance of animals was then tested on previously learnt odor and airflow rate discrimination tasks. While the performance of treated mice dropped to chance level, control mice performed with an accuracy of >80% in a previously learned odor discrimination task, thereby confirming the ablation of OSNs. On repeating the training with airflow rate discriminations, the control group performed with high accuracy while the treated animals performed at chance levels (Figure 2D1, two-way ANOVA with Tukey’s multiple comparison, * represent p < 0.05). The ITIs, measure of animals’ motivation, remained optimal and unaltered even after the treatments (Figure 2D2, one-way ANOVA for individual airflow/odor pairs, p > 0.05). These results reveal the critical role of OSNs in anemo-discrimination.

**Figure 2.**
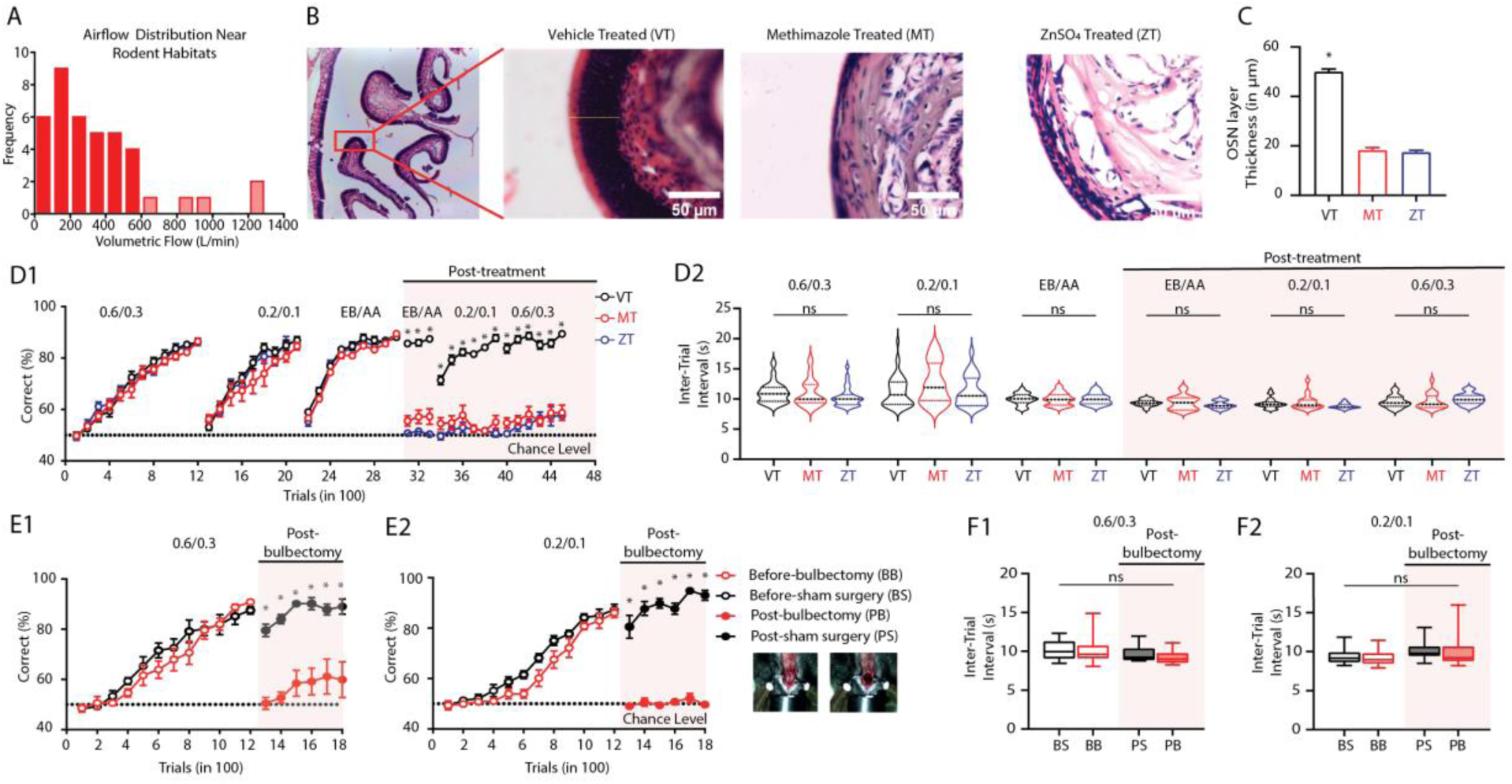
Mice use olfactory system to detect and discriminate airflows. **A.** Histogram representing the distribution of airflows in potential rodent habitats. The airflows in the potential rodent habitats were measured using a hot-wire anemometer from 40 different locations. 90 % of airflow rates that were observed in the potential habitats were in between 0.1 and 0.6 LPM (shown by solid red bars), with 0.35 LPM being the average airflow rate. **B.** Images showing Hematoxylin and Eosin-stained section of olfactory epithelium under different treatment conditions. **C.** Thickness of epithelium measured across vehicle, methimazole and zinc sulphate administered animals after two days of the treatment. Epithelium thickness for vehicle (VT: Vehicle Treated) was significantly higher as compared to the width of epithelium in methimazole (MT) and zinc sulphate (ZT) treated animals (one-way ANOVA, F = 354.2, p<0.0001, n =2 mice for all groups, N = 50 region of interests from 6 sections from each mouse). **D. D1.** Learning curve representing discriminating accuracies of animals for different airflow and odor pairs before and after the treatment. The learning accuracies of animals that underwent ZT and MT were significantly lower to that of the control group. Statistical comparison across different groups for each airflow or odor pair were done by using two-way ANOVA with Tukey’s multiple comparison. * represents p<0.05. (n_VT_ = 14-15, n_MT_ = 10, n_ZT_ = 8). **D2.** Violin plots representing inter-trial interval (ITI) of animals, which is used as a readout for motivation levels. The ITI is compared for animals across different groups across different airflow and odor pairs under different OSN ablation treatments. The ITI of all animals throughout the training remained in the range of 9-11 s. Statistical significance was measured using one-way ANOVA with Tukey’s multiple comparison and p-value of > 0.05 was observed across different groups for different airflow and odor pairs. **E. E1.** Learning of two groups of animals for 0.6 vs 0.3 LPM before and after the bulbectomy surgery. Before the surgery both bulbectomized and sham surgery group showed similar learning pace (two-way ANOVA, F(11,132) = 1.282, p = 0.2415, n_bulbectomy_ = 8, n_sham_ = 6). Following surgery, the performance of bulbectomized group dropped to the chance level and remained so for 600 trials, however, the sham surgery group did not show any performance deficit (two-way ANOVA with Tukey’s multiple comparison test, * represents p<0.05, n_bulbectomy_ = 7, n_sham_ = 6). **E2.** Learning of two groups of animals for 0.2 vs 0.1 LPM before and after the surgery. Before the surgery both bulbectomized and sham surgery group showed similar learning pace (two-way ANOVA, F(11,132) = 1.162, p = 0.3198, n_bulbectomy_ = 8, n_sham_ = 6). Following surgery, the performance of bulbectomized group dropped to the chance level and remained so for 600 trials, however, the sham surgery group did not show any performance deficit (two-way ANOVA with Tukey’s multiple comparison test, * represents p<0.05, n_bulbectomy_ = 5, n_sham_ = 6). **F. F1.** Inter-trial intervals of different groups of animals for 0.6 vs. 0.3 LPM before and after the surgery. ITI of the animals did not reveal any significant differences across different groups and surgical manipulations (one-way ANOVA, F = 1.841, p = 0.1466). **F2.** Inter-trial intervals of different group of animals for 0.2 vs. 0.1 LPM before and after the surgery. ITI of the animals did not reveal any significant differences across different groups and surgical manipulations (one-way ANOVA, F = 2.304, p = 0.0839).

Once we confirmed the olfactory route of airflow sensation, we then studied the role of OB, the first relay station in olfactory signaling, in processing airflow rate related information. Mice’ airflow discrimination abilities were measured before and after surgically removing the OB. Two groups of experimental mice and corresponding control animals (sham surgery) were trained to discriminate 0.2 vs. 0.1 LPM and 0.6 vs. 0.3 LPM Animals performed with > 80% accuracy within 1200 trials for both the airflow pairs. Experimental and control mice did not show any difference in learning pace before the treatment (Figure 2E1, 2E2, two-way ANOVA with Tukey’s multiple comparison, p >0.05 for both groups). Once the animals reached the asymptotic phase of learning, OB was surgically aspirated and animals were allowed a recovery period of 12-15 days. Training was then continued with all mice to discriminate the already learned airflow pair. Post bulbectomy, the performance of experimental mice dropped to chance levels for both airflow pairs, where no alteration in the performance was observed for the control animals that underwent sham surgery (Figure 2E1, 2E2, two-way ANOVA with Tukey’s multiple comparison, * represent p < 0.05). We did not observe any differences in the ITIs for all groups of animals for both airflow pairs, indicating that the motivational levels of animals were optimal and comparable before and after the treatment (Figure 2F1, 2F2, one-way ANOVA, p>0.05). These findings confirm the involvement of OB in anemo-discriminations.

### Stimulus-independent and learning-dependent refinement of sniffing during the decision-making period in airflow rate discrimination

As we observed the loss of airflow rate discrimination abilities on blocking the airflow information processing through olfactory pathway, we next probed modulation in their sampling behavior during airflow rate discriminations. We trained another batch of mice under head-restrained conditions and monitored their sniffing behavior (see Methods), an exploratory behavior associated with olfaction, during different phases of learning (Bhattacharjee et al., 2019). Mice were trained on different airflow rates until they reached the performance accuracy of >80% (0.2 vs. 0.1 LPM [1200 trials], 0.6 vs. 0.3 LPM [900 trials] and 0.4 vs. 0.45 LPM [900 trials], Figure 3A). As animals learned to discriminate the first airflow pair with high accuracy, which was also reflected by faster FDTs, we observed significant changes in breathing modulations, quantified by breath initiation counts (BICs) and other sniffing associated parameters (sniffing frequency (SF), Figure 3A-E). The raster plots of breath initiations and the related histograms showed that at the early stages of learning (First task), BICs remained higher even after the sniffing peak, however, at the learned phase (Last task) BICs were comparable to that of pre-stimulus duration (Figure 3B1, B2). This was further reflected in the SF measurements during the entire stimulus duration and the decision-making period which is the same as discrimination time. While SF during stimulus duration decreased from first to last task, SFs during the decision-making period showed a significant enhancement with learning (Figure 3C, D, two-tailed paired t-test, *represents p<0.05). However, the inhalation onset remained unchanged and was independent of learning phases (Figure 3E, two-tailed paired t-test, p = 0.1955). Further, we analyzed SFs before-, after- and during-FDTs when animals were performing with >80% accuracy for all three airflow pairs they were trained on. Mice showed higher sniffing frequencies during the FDTs compared to before and after the decision-making duration and were independent of reward contingencies as well (Figure 3F, Supplementary Figure S3). The SFs during the entire stimulus duration, SFs during the decision-making period, and inhalation onsets remained similar for all airflow pairs tested (Figure 3G, 3H and 3I, one-way repeated measures ANOVA, p>0.05). These results provide conclusive evidence for stimulus-independent and learning-dependent temporal refinement of sniffing confined to decision-making period, which may help animals to take accurate decisions.

**Figure 3.**
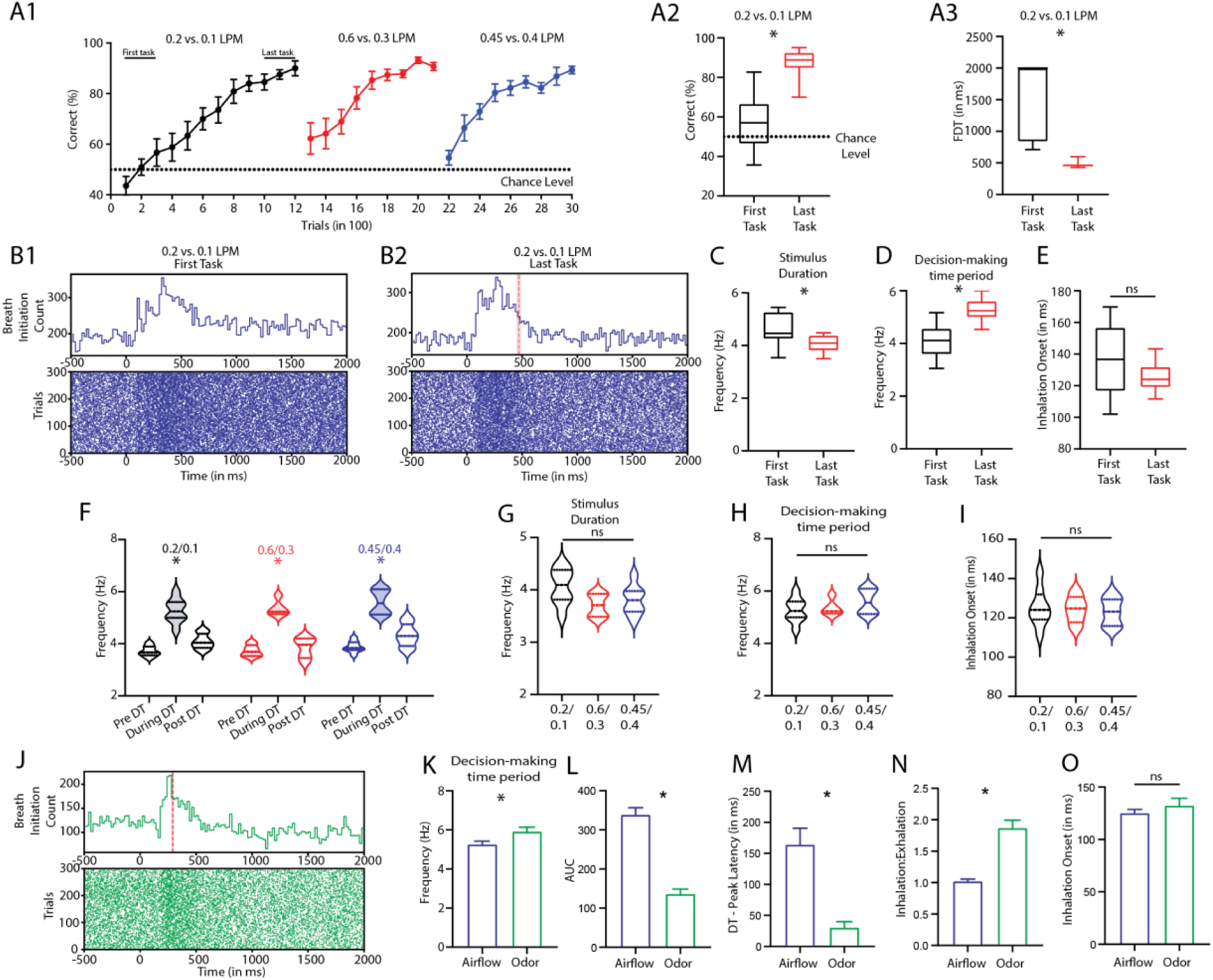
Mice exhibit learning-dependent and stimulus-independent sniffing refinement during anemo-discrimination. **A. A1.** Learning curves of animals trained on different airflow pairs (0.2 vs. 0.1 LPM, 0.6 vs. 0.3 LPM and 0.45 vs. 0.4 LPM). **A2.** Box and whisker plots showing accuracies of animals during first and last task of the training for 0.2 vs. 0.1 LPM. The accuracy of animals was significantly higher for last task (two-tailed paired t-test, p = 0.0012, n = 8). **A3.** Box and whisker plots showing FDTs of animals during first and last task of the training for 0.2 vs. 0.1 LPM. The FDT of animals was significantly lower for the last task (two-tailed paired t-test, p = 0.0013, n = 8). **B. B1. and B2.** Raster plots and histogram representing inhalation onset during first and last task while animals were trained to discriminate 0.2 vs. 0.1 LPM. **C.** Box and whisker plots showing sniffing frequency of animals during 2s stimulus duration for first and last task of the training for 0.2 vs. 0.1 LPM. The SF of animals was significantly lower for the last task (two-tailed paired t-test, p = 0.0157, n = 8). **D.** Box and whisker plots showing SF of animals during decision-making window for first and last task of the training for 0.2 vs. 0.1 LPM. The SF of animals during the decision-making window was significantly higher for the last task (two-tailed paired t-test, p = 0.0009, n = 8). **E.** Box and whisker plots showing onset of first inhalation during decision-making window for first and last task of the training. The inhalation onset for first and last task was similar (two-tailed paired t-test, p = 0.1955, n = 8). **F.** Violin plots showing sniff frequencies of animals for before, during and post decision-making period across different airflow pairs. Animals showed higher SF during decision-making period across different airflow pairs (0.2 vs. 0.1 LPM; one-way repeated measures ANOVA, F = 141.3, p < 0.0001, n = 8, 0.6 vs. 0.3 LPM; one-way repeated measures ANOVA with Tukey’s multiple comparison test, F = 54.42, p = 0.0005, n = 5, 0.45 vs. 0.4 LPM; one-way repeated measures ANOVA, F = 139.5, p < 0.0001, n = 8). **G.** Violin plots showing sniffing frequencies of animals during 2 s stimulus period for different airflow pairs. Animals showed similar SFs for different airflow pairs (one-way repeated measures ANOVA, F = 2.662, p = 0.0971, n = 5-8). **H.** Violin plots showing sniff frequencies of animals during decision-making period across different airflow pairs. The SFs of animals during the decision-making period were similar across different airflow pairs (one-way repeated measures ANOVA, F = 1.211, p = 0.3211, n = 5-8). **I.** Violin plots showing onset of first inhalation during decision-making window across different airflow pairs. The inhalation onset of animals for different airflow pairs were non-significant (one-way repeated measures ANOVA, F = 0.1481, p = 0.8634, n = 5-8). **J.** Raster plots and histogram representing inhalation onset for odor-based discrimination (Binary mixture of enantiomers [Octanol (+) vs. Octanol (–)]) pooled across animals in the last phase of learning (last 300 trials). The red vertical line represents the odor discrimination time (ODT) and the shaded region represents SEM. **K.** Bar graphs representing sniff frequencies of animals for airflow (0.2 vs. 0.1 LPM) and odor trials during decision-making period. The SFs of animals for odors were significantly higher (SF: airflow discrimination – 5.268 ± 0.1635, odor discrimination – 5.925 ± 0.2156, two-tailed unpaired t-test, p = 0.0291, n_airflow_ = 8, n_odors_ = 6). **L.** Bar graphs representing area under the curve (AUC) for airflow and odor trials during decision-making period. The AUC is calculated for sniff histograms from individual animals and averaged. The AUC for airflows was significantly higher (AUC: airflow discrimination – 339.1 ± 17.43, odor discrimination – 137.0 ± 11.65, two-tailed unpaired t-test, p < 0.0001, n_airflow_ = 8, n_odors_ = 6). **M.** Bar graphs representing difference in the FDT and peak latency for airflow and odor trials during decision-making period. The difference is calculated for individual animals and averaged. The difference was significantly higher for airflows (airflow discrimination – 164.3 ± 26.35 ms, odors – 30.67 ± 9.262 ms, two-tailed unpaired t-test, p = 0.0012, n_airflow_ = 8, n_odors_ = 6). **N.** Bar graphs representing ratio of time spent in inhalation to that of the time spent in exhalation for airflow and odor trials during decision-making period. The ratio of inhalation to exhalation time was significantly lower for airflows (airflows – 1.024 ± 0.0304, odors – 1.871 ± 0.1229, two-tailed unpaired t-test, p < 0.0001, n_airflow_ = 8, n_odors_ = 6). **O.** Bar graphs representing onset of first inhalation for airflow and odor trials during decision-making period. Animals showed similar inhalation onset for airflows and odors (Inhalation Onset: airflow discrimination – 125.3 ± 3.424 ms, odor discrimination – 132.4 ± 6.972 ms, two-tailed unpaired t-test, p = 0.3419, n_airflow_ = 8, n_odors_ = 6).

### Distinct sampling behaviors are shown by mice for airflow- and odor discrimination

Rodents are known to modulate their sniffing behavior in odor discrimination contexts (Bhattacharjee et al., 2019; Wachowiak, 2011). As we observed significant sniffing modulations shown by mice during airflow discriminations, we further compared the sampling behavior during odor and airflow discriminations. We trained a new group of animals on an odor discrimination task in which they had to distinguish between a binary mixture of enantiomers [Octanol (+) vs. Octanol (–)]. After the animals reached a performance accuracy of >80%, we quantified various sniffing parameters and compared them with that of airflow discriminations. Mice showed distinct temporal patterns of breathing during airflow and odor discrimination behavior (Figure 3B2, J). The sniffing frequency of animals increased for both airflows and odors during the decision-making window, but the sniffing frequencies for odors were significantly greater than that for airflows (Figure 3K, two-tailed unpaired t-test, p = 0.0291). Further, breath initiation histograms revealed that the animals displayed enhanced sniffing for longer durations in case of airflow discrimination compared to odor discrimination. This was quantified by the area under the curve (AUC) of the histogram that was generated by plotting breath initiations on individual trial basis and averaging across animals (Figure 3L, two-tailed unpaired t-test, p < 0.0001).

As we observed different discrimination time for odor and airflow discriminations, we calculated the difference between sniffing peak latency and discrimination time (DT-peak latency). The difference was greater for airflow discriminations compared to odor discriminations, implying that animals need longer time to make airflow rate-based decisions at optimal sniff frequencies than odorants (Figure 3M, two-tailed unpaired t-test, p = 0.0012). Further, to understand breathing dynamics in detail, we analysed the inhalation-exhalation times shown by animals. During odor discrimination, we observed a significantly higher inhalation-exhalation time ratio than airflow discrimination (Figure 3N, two-tailed unpaired t-test, p < 0.0001). This may be due to animals’ attempt to collect maximum odor information in a discrimination context. Despite the changes in several sniff parameters, the time of first inhalation remained similar for both the odor and airflow discriminations (Figure 3O, two-tailed unpaired t-test, p = 0.3419), implying that first inhalation happens independently of stimulus characteristics and the temporal dynamics of breathing evolves depending on the kind of stimulus. Together, these results confirm the involvement of mouse olfactory system in anemo-discrimination and prove that animals employ distinct sniffing strategies in gathering sensory information while challenged with airflow rate and odor discriminations.

### Olfactory bulb inhibitory network regulates anemo-detection and -discrimination

As surgical removal of OB led to inability of mice to execute anemo-processing tasks, we then dissected the underlying circuitry in OB-intact mice. To do this, the neuronal activation pattern in OB following the airflow discrimination task, were examined using c-Fos. Neurons in different layers of OB expressed c-Fos, whereas the signal was most abundant in granule cell layer (GCL) that harbors most of the GAD65 (GAD2) positive (an isoform of GABA synthesizing enzyme) GABAergic inhibitory interneurons (EYFP labeled, Figure 4A). We further studied how OB inhibitory network responds towards airflow stimuli by quantifying Ca^2+^ dynamics using specific expression of GCaMP6f in GAD65 interneurons. Grin-lenses were implanted in the GCL of GAD65-GCaMP6f transgenic mice, and calcium dynamics were monitored while they were presented with various airflow rates (see Methods and Figure 4B). The population activity of OB GAD65 interneurons were investigated under both anaesthetized and awake conditions. While stimulus strength dependent activation was studied under anesthesia, effect of airflow discrimination learning on the interneuronal population activity was monitored during different phases of learning. On quantifying fluorescence changes in response to different airflows under anesthetized conditions, the DF/F0 increased after the stimulus onset, reached a peak and fell back to the baseline (Figure 4C1). The fluorescence changes were quantified for 250 trials across 5 animals for various airflow rates (Figure 4C2). On averaging DF/F0 across all animals, we observed the increase in amplitudes correlating with stimulus strength (Figure 4D, Pearson correlation: R^2^ = 0.9911, p = 0.0045). This confirms the involvement of OB GABAergic interneurons in processing airflow-related information.

**Figure 4.**
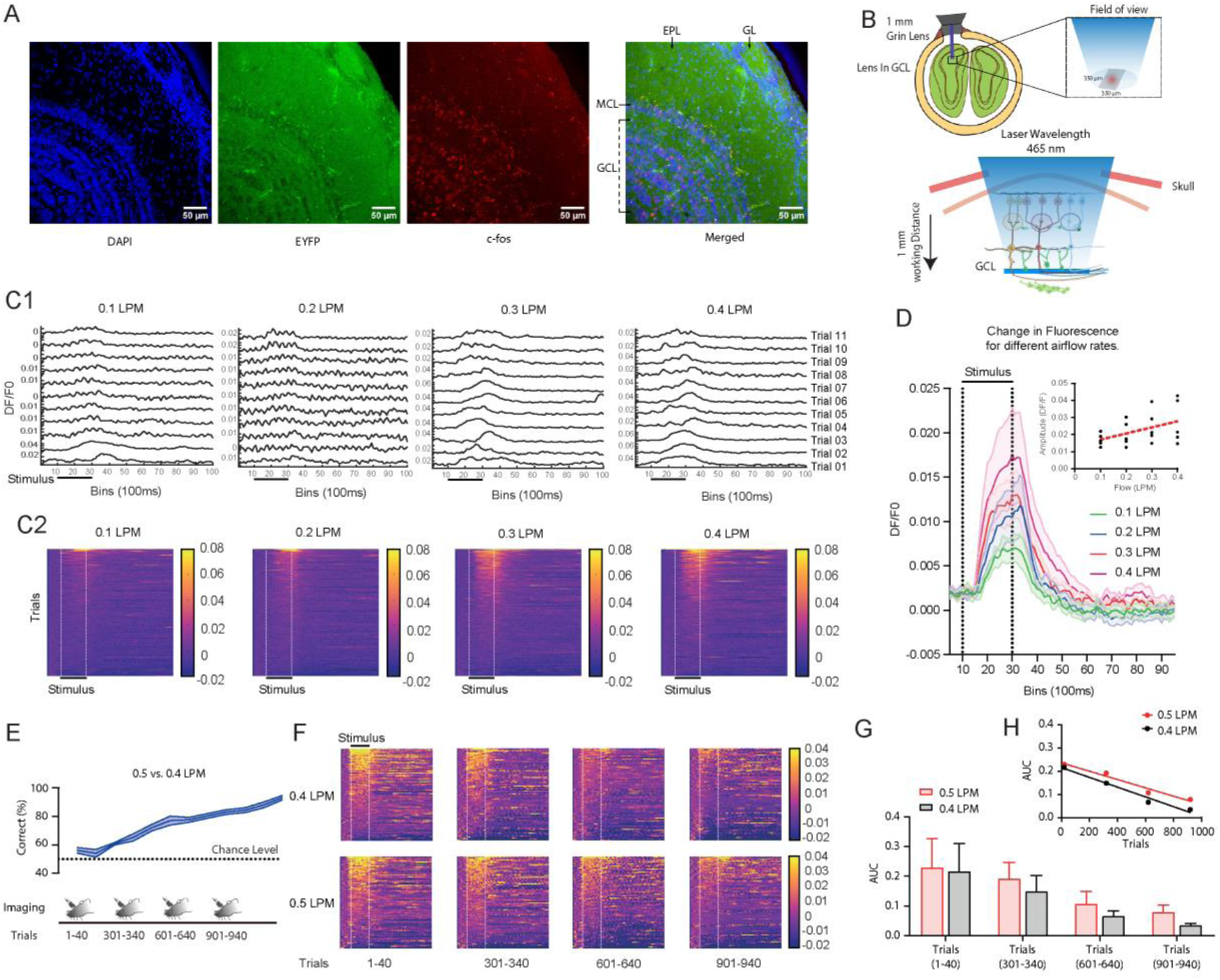
GABAergic (GAD65+ve) OB interneurons are involved in processing airflow stimuli in a learning-dependent manner. **A.** c-Fos expression in various OB layers of animals trained to distinguish different airflows. The GCL layer, which houses the majority of the GAD65-expressing interneurons, has the highest level of c-Fos expression. Blue - DAPI staining, Green – GAD65 - EYFP expression, Red - c-Fos expression. **B.** Illustration representing the scheme of micro-endoscopic calcium imaging. Depth range and field of view of imaging cannula used for imaging GCs is displayed. **C. C1.** Trial-by-trial traces of GAD65 +ve OB interneuron response to different airflows in a single animal under anaesthetized conditions. **C2.** Pseudo-color heatmaps of Ca^2+^ dynamic responses elicited by various airflows. Data were collected from 5 mice, with 20 trials for each airflow. Data from all trials are sorted in decreasing order of Ca^2+^ responses during airflow administration. For all the airflows, GAD65+ve interneurons showed activity towards airflows during the stimulus presentation period. **D.** The average DF/F0 responses of all animals for various airflows. The solid line represents the mean across all animals, while the shaded area represents the standard error of the mean (SEM). The DF/F0 increased after the stimulus presentation and decayed after the stimulus offset. The subgraph depicts the change in amplitude as the stimulus strength increases. Individual dots represent animal wise amplitude values. The amplitude is found to be positively correlated with stimulus strength (Pearson correlation: R^2^ = 0.9911, p = 0.0045, n = 5 mice). **E.** The learning curve represents accuracy of performance as the training progresses. The solid line shows the average accuracy of all animals, while the shaded area shows the standard error of the mean. Additionally, schematic for the sequence of imaging sessions of the experiment is provided. The imaging is done at the beginning of each task for 40 trials. **F.** Pseudo-color heatmaps of Ca^2+^ responses elicited by 0.4 and 0.5 LPM for different tasks. Data were collected from 5-6 mice, with 20 trials for each airflow at the beginning of each task. The qualitative analysis of heatmaps show a potential refinement in the Ca^2+^ transients with learning. Data from all trials are sorted in decreasing order of Ca^2+^ response during the presentation of airflow stimulus. **G.** Bar graphs representing average AUC of animals for 0.4 and 0.5 LPM with progression of trials. As learning progressed, the AUC for both airflows decreased. **H.** Plot representing change in average AUC of animals for 0.4 and 0.5 LPM with number of trials. For both airflows, change in average AUC showed an inverse correlation with learning (0.4 LPM: R^2^ = 0.9726, p = 0.0138; 0.5 LPM: R^2^ = 0.9636, p = 0.0184, n = 5-6 mice).

Having observed the involvement of inhibitory network in airflow information processing, we further investigated the modulation of inhibition in different learning phases. A group of mice that express GCaMP6f in GAD65 interneurons, were trained to discriminate the airflows 0.5 vs. 0.4 LPM (1200 trials, 4 tasks) under head-restrained conditions. The calcium activity was recorded from the first 40 trials of each task (300 trials) from each animal. Animals learned to discriminate this flow pair with a high accuracy of >80% (Figure 4E). We observed calcium activity in response to both rewarded and non-rewarded stimuli throughout the discrimination training duration, however, in a decreasing manner as the learning progressed (Figure 4E, F). To quantify this Ca^2+^ dynamics, the area under the curves (AUCs) from the DF/F0 traces for both rewarded and non-rewarded stimuli were calculated. A decrease in AUCs were observed for both stimuli as the accuracy of discrimination performance improved from chance level to ∼90% correct responses (Figure 4G, H, 0.4 LPM: R^2^ = 0.9726, p = 0.0138; 0.5 LPM: R^2^ = 0.9636, p = 0.0184). These findings provide the first experimental evidence for the modulation of OB inhibitory interneuronal population activity during the airflow discrimination in a learning-dependent fashion.

### Synaptic inhibition in the mouse olfactory bulb controls anemo-discrimination behavior

Having observed the learning-dependent refinement of calcium activity in the inhibitory neurons during airflow discriminations, we further tried to probe how the modulation of these neurons affect anemo-discriminations. We started by studying the anemo-discrimination behavior shown by knockout mice where the AMPA receptor subunit, GluA2 deletion was specific to GAD65 interneurons (Figure 5A, B). Knocking out GluA2 subunit from inhibitory granule cells of OB resulted in the enhancement of Ca^2+^ influx and synaptic inhibition which facilitated odor discrimination (Abraham et al., 2010). We trained one batch of mice to discriminate 0.5 vs. 0.4 LPM and we observed slower learning pace with the GluA2 KOs compared to control mice (Figure 5C, two-way ANOVA, *represents p<0.05). This observation was in contrast to the effect of synaptic inhibition on odor discrimination behavior, which indicates that the extent of inhibition needed to refine the airflow information is different from that of the odor information. Therefore, we decided to characterize the phenotypes resulted by bidirectional optogenetic OB-specific modulation of GAD65 inhibitory interneurons.

**Figure 5.**
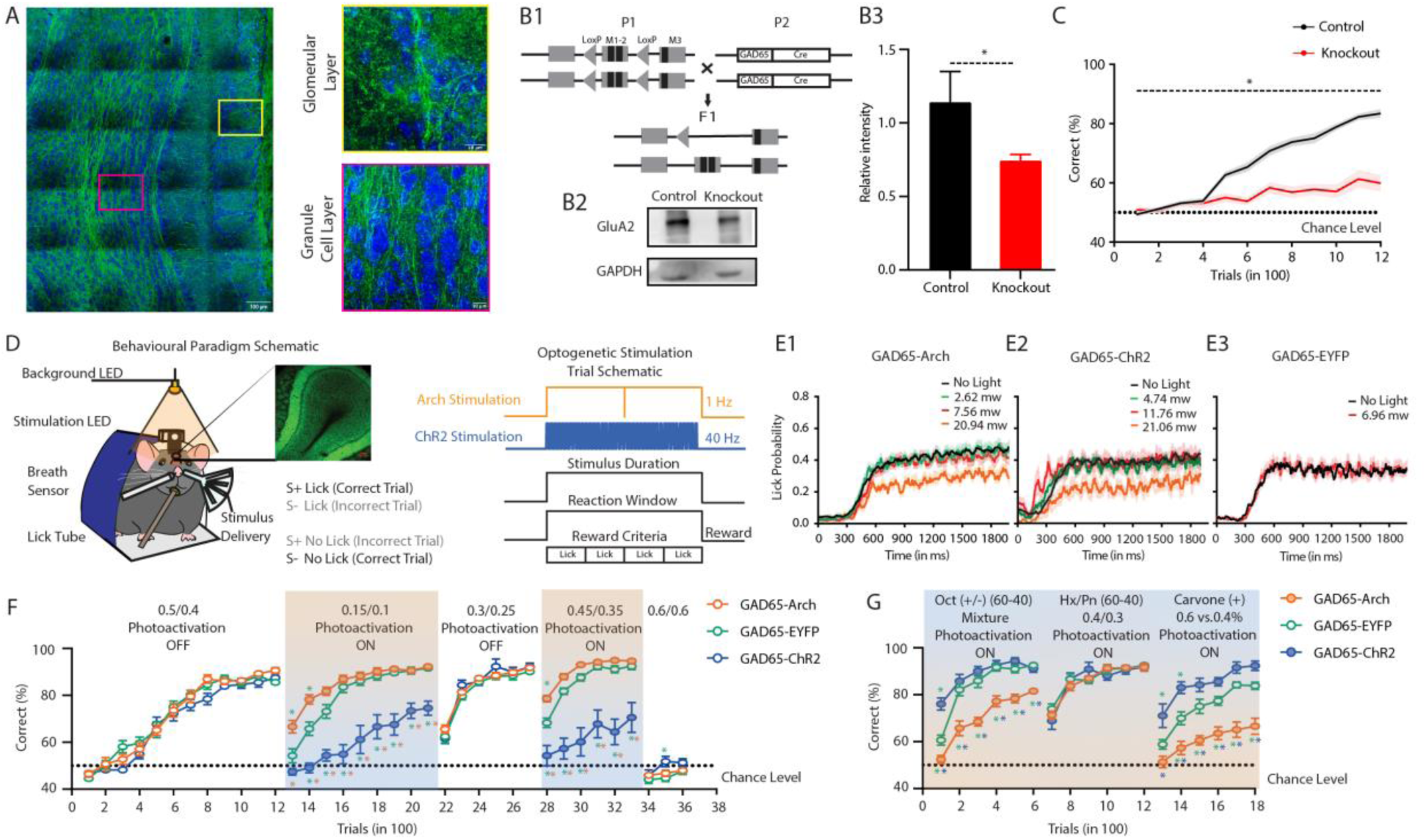
Synaptic inhibition in the mouse olfactory bulb controls anemo-detection and discrimination. **A.** Tile scan representing the expression of GAD65 (GAD2) in the olfactory bulb of mice. Enlarged images (towards the right) show the GAD65 expression in the different layers of the olfactory bulb. Green represents GAD65 expression whereas blue represents DAPI. **B. B1**. Scheme of genetic crosses used to create the GAD65-GluA2 knockout animals. **B2.** Western blot showing the expression of GluA2 protein in the olfactory bulb of controls as well as GAD65-GluA2 knockout animals. Band corresponding to GAPDH protein is observed in olfactory bulb lysates from both animals. **B3.** Relative GluA2 expression in control animals (measured by normalizing the GluA2 intensity to that of corresponding GAPDH). GluA2 expression is reduced significantly in knockout animals (two-tailed unpaired t-test, p = 0.0347, n_Controls_ = 5, n_GAD65-GluA2 knockout_ = 6). **C.** Comparison of learning curves for control and GAD65-GluA2 knockout animals trained to discriminate 0.5 vs. 0.4 LPM. GAD65-GluA2 knockout animals showed significant lower learning accuracy as compare to that of control animals (Two-way ANOVA, F = 298.6, p<0.0001, n_control_ = 18, n_GAD65-GluA2 knockout_ = 16). **D. Left.** Experimental setup to perform behavioral training under head-restrained conditions. Animal with implanted LED is head-restrained on a metallic plate. The stimulus is delivered through a tube and LED stimulation takes place during the stimulus presentation. During the stimulus presentation animal has to lick on the lick tube to get the water reward. The breathing is also monitored during the training. **Right.** Illustration representing optogenetic (LED) stimulation. A 595 nm amber LED is fixed on top of a circular cranial window over the OB in a GAD65-Arch expressing mouse and a 473 nm blue LED is fixed on top of a circular cranial window over the OB in a GAD65-ChR2 expressing mouse. The response window, wherein animal has to lick to get the reward coincide with the stimulus delivery period. **E.** LED power optimization for optogenetic modification of GAD65 +ve interneurons during an airflow detection task. **E1.** Average lick responses of GAD65-Arch mice to different light intensities. The solid lines and shaded areas represent the mean and standard error of mean, respectively. Because each mouse licked differentially at different LED powers, the LED power for further discrimination experiments was determined individually. **E2.** Average lick responses of GAD65-ChR2 mice to different light intensities. The solid lines and shaded areas represent the mean and standard error of mean, respectively. Because each mouse licked differently at different LED powers, the LED power was determined individually. **E3.** Average lick responses of GAD65-EYFP mice at no light and light at 6.96 mW (which is the average light intensity used for further experiments for ChR2 animals). The solid lines and shaded areas represent the mean and standard error of mean, respectively. **F.** Learning curves representing performance accuracy for various airflow pairs under different photoactivation conditions. Animals were trained in the following order: 0.5 vs. 0.4 LPM (1200 trials - Light OFF), 0.15 vs. 0.1 LPM (900 trials - Light ON), 0.3 vs. 0.25 LPM (600 trials - Light OFF), 0.45 vs. 0.35 LPM (600 trials - Light ON), 0.6 vs. 0.6 LPM (600 trials - Light ON). Animals learnt faster when GAD65-expressing interneurons were stimulated with Archaerhodopsin, whereas a deficit in learning occurred for Channelrhodopsin activation (Two-way ANOVA with Tukey’s multiple comparison test: * represents p< 0.05). **G.** Learning curves representing animal accuracy for various odor-based tasks using same/different airflow rates. Animals were trained in the following order: Octanol (+) vs. Octanol (–) - 60-40 mixture (600 trials - Light ON), Hexanal/Pentanone - 60-40 mixture - 0.4/0.3 LPM (600 trials - Light ON), Carvone (+) 0.6 vs. 0.4 % (v/v, 600 trials - Light ON), Two-way ANOVA with Tukey’s multiple comparisons. (* represents p< 0.05).

The experimental animals consisted of different groups of mice expressing Channelrhodopsin (ChR2) and Archeorhodopsin (Arch) in the GAD65 interneurons of the OB, whereas control animals had EYFP expressed in these interneurons. OB-specific modulation was achieved by implanting the LED on top of the OB (Figure 5D, See STAR Methods). To identify the optimal intensity of photoactivation, we studied the airflow detection behavior of mice in presence of varying stimulation light intensities. For both ChR2 and Arch expressing animals, the licking responses decreased as the intensity of photo-activation increased (Figure 5E1, E2), whereas no change in the licking responses of control animals were observed. For further experiments, the light intensities were chosen in such a way that the licking responses of the animals during photoactivation remained similar to that in the no-photoactivation condition [Arch - range: 1-5 mW, mean: 2.93 mW), ChR2 - range: 3-11 mW, mean: 6.96 mW, and EYFP - mean: 6.96 mW (same for all animals)].

Further, the role of inhibitory circuitry in controlling airflow discrimination efficiency was examined. Different groups of animals were trained on a sequence of airflow discrimination tasks wherein photo-activation ON and OFF conditions were intermittently applied. For the first airflow pair (0.5 vs. 0.4 LPM - Light OFF) animals reached the high-accuracy, asymptotic phase within 1200 trials. There was no difference in the learning pace across different groups, suggesting that intrinsic differences do not exist among the groups that can influence learning efficiency. However, when the animals were trained on 0.15 vs. 0.1 LPM under photo-activation conditions, we observed faster learning when GAD65 expressing interneurons were inhibited by Arch stimulation and slower learning during ChR2 stimulation (Figure 5F, two-way ANOVA with Tukey’s multiple comparison test, * represents p<0.05). A similar trend was observed for another airflow pair 0.45 vs. 0.35 LPM during the photo-activation conditions. The observed bidirectional modulations in the learning pace were specific to photo-activation conditions, as the animals did not show any difference in their learning pace while trained on another airflow pair (0.25 vs. 0.3 LPM) without any photo-activation (Figure 5F, two-way ANOVA with Tukey’s multiple comparison test, * represents p<0.05). On training the mice with same airflow rate (0.6 vs. 0.6 LPM), animals’ performance remained at chance levels confirming the specific learning of airflow discriminations (Figure 5F). As a result of photoactivation, the FDTs were also altered. ChR2-expressing animals showed slower FDTs when compared across different groups (Supplementary Figure S4A1). For all the animals performing above 80% across different airflow pairs under light-ON conditions, the Arch group showed significant improvement in FDTs as compared to control groups (Supplementary Figure S5). However, as the ChR2-expressing animals’ average performance accuracy was below 80%, this analysis could not be performed. These phenotypic differences did not arise as a result of varying sampling strategies that may occur due to optogenetic stimulations. In all groups, animals showed similar sniffing frequencies across different airflow pairs (Supplementary Figure S4B1), indicating that the alterations in airflow discrimination behavior was specifically due to the modulations of OB inhibitory circuitry.

We continued to study the role of OB inhibitory circuitry in controlling odor discrimination. In contrast to airflow discriminations, opposing phenotypes were observed when the animals were trained on an odor discrimination task (Figure 5G, Octanol complex mixture: 60-40 vs. 40-60 binary mixtures). The phenotype observed was consistent with previous findings where enhanced inhibition led to better odor discrimination abilities(Abraham et al., 2010; Gschwend et al., 2015). The ODT shown by Arch group was significantly slower, and no difference in the sniffing frequencies were observed between groups (Supplementary Figure S4A2, B2). As contrasting phenotypes were observed for airflow- and odor-based discrimination tasks, we studied the changes in multimodal stimuli discrimination efficiency (Odor A + Airflow 1 vs. Odor B + Airflow 2, 60-40 Hexanal-Pentanone + 0.4 LPM vs. 40-60 Hexanal-Pentanone + 0.3 LPM) upon optogenetic modulation. All three groups of mice learned this discrimination with similar efficiencies. This indicates that the deficiencies we observed either for odor- or airflow-based discriminations under optogenetic modulations were rescued by altering the second modality added to the multimodal stimuli (Figure 5G, two-way ANOVA with Tukey’s multiple comparison test, * represents p<0.05). Similar DTs were observed for different groups, however, an increase in sniffing frequency was observed for ChR2 group (Supplementary Figure S4A2, B2). Further experiments dissecting the connectivity between breathing and learning centers are needed to find out the reasons for this modulation.

Varying airflows associated with specific stimuli can cause concentration differences of odorants. Therefore, we varied the airflows with a specific stimulus and challenged the animals to discriminate while bidirectionally modulating the inhibitory network activity. The phenotype during this concentration-based discrimination task was similar to that observed during the odor-based task; ChR2 group learnt faster, whereas Arch group showed learning deficits and higher ODT (Figure 5G, two-way ANOVA with Tukey’s multiple comparison test, * represents p<0.05). All groups of mice showed similar sniffing frequencies during this task, while ODT for Arch animals were found to be slower (Supplementary Figure S4A2, B2). Taken together, the bi-directional changes of behavioral readouts during the airflow and odor discrimination tasks, and the rescue of the learning deficits during multimodal tasks under optogenetic modulations, imply that synaptic inhibition in olfactory bulb refines the airflow and odor information processing. However, the optimal inhibition required for airflow- and odor-information refinement varies. Moreover, OB neural circuits might be playing a critical role in integrating the chemical and mechanical information and generating the ‘multimodal’ odor percept in the dynamic olfactory world.

### Mechanosensation through mouse nose enhances olfactory perception at subthreshold stimuli conditions

In nature, rodents sample and perceive odorants carried by turbulent airflows. These odor plumes and the associated airflows make the olfactory environment very dynamic and may challenge animals to detect and perceive chemical cues even at the subthreshold concentrations. Here, we tried to address the functional relevance of mechanosensation through rodent nose. To test our hypothesis of mechanosensation aiding olfactory perception at subthreshold conditions, we started training different batches of mice on varying odor concentrations and airflow rates. To find out the minimum differences in airflow rates that mice can discriminate, they were trained on different airflow rates and three flow pairs were concluded as below the discrimination threshold - 0.1 vs. 0.11 LPM, 0.2 vs. 0.23 LPM, and 0.3 vs. 0.34 LPM (Supplementary Figure S6A). These discrimination thresholds were following Weber’s law (Pardo-Vazquez et al., 2019; Weber, 1834)(Supplementary Figure S6B, C). Further, the discrimination thresholds were calculated for (+) Carvone vs. (–) Carvone (0.01 % v/v), and Cineol vs. Eugenol (0.01 % v/v, Supplementary Figure S7). Next, we trained three different groups of mice on subthreshold discrimination levels for airflow rates, odors and on the ‘multimodal’ stimuli combining both airflow rates and odors i.e. (+) Carvone vs. (–) Carvone (at 0.3 vs. 0.34 LPM), and Cineol vs. Eugenol (at 0.2 vs. 0.23 LPM). Compared to odor-only and airflow-only groups, mice that were trained on ‘multimodal’ stimuli showed significantly enhanced learning pace (Figure 6A1, A2). Further, to probe if sampling differences could account for the phenotypes we observed, another batch of mice were trained on similar stimuli under head-restrained conditions and their sniffing behavior was quantified (Figure 6B). The ‘multimodal’ group showed high-performance levels of >80% at the end of training compared to other groups. On quantifying the sniffing frequency and inhalation onsets, we observed enhanced breathing under multimodal conditions compared to unimodal ones (Figure 6C1, C2). These results provide robust evidence for the facilitation of olfactory discrimination by mechanosensation through the mouse olfactory system, at subthreshold levels.

**Figure 6.**
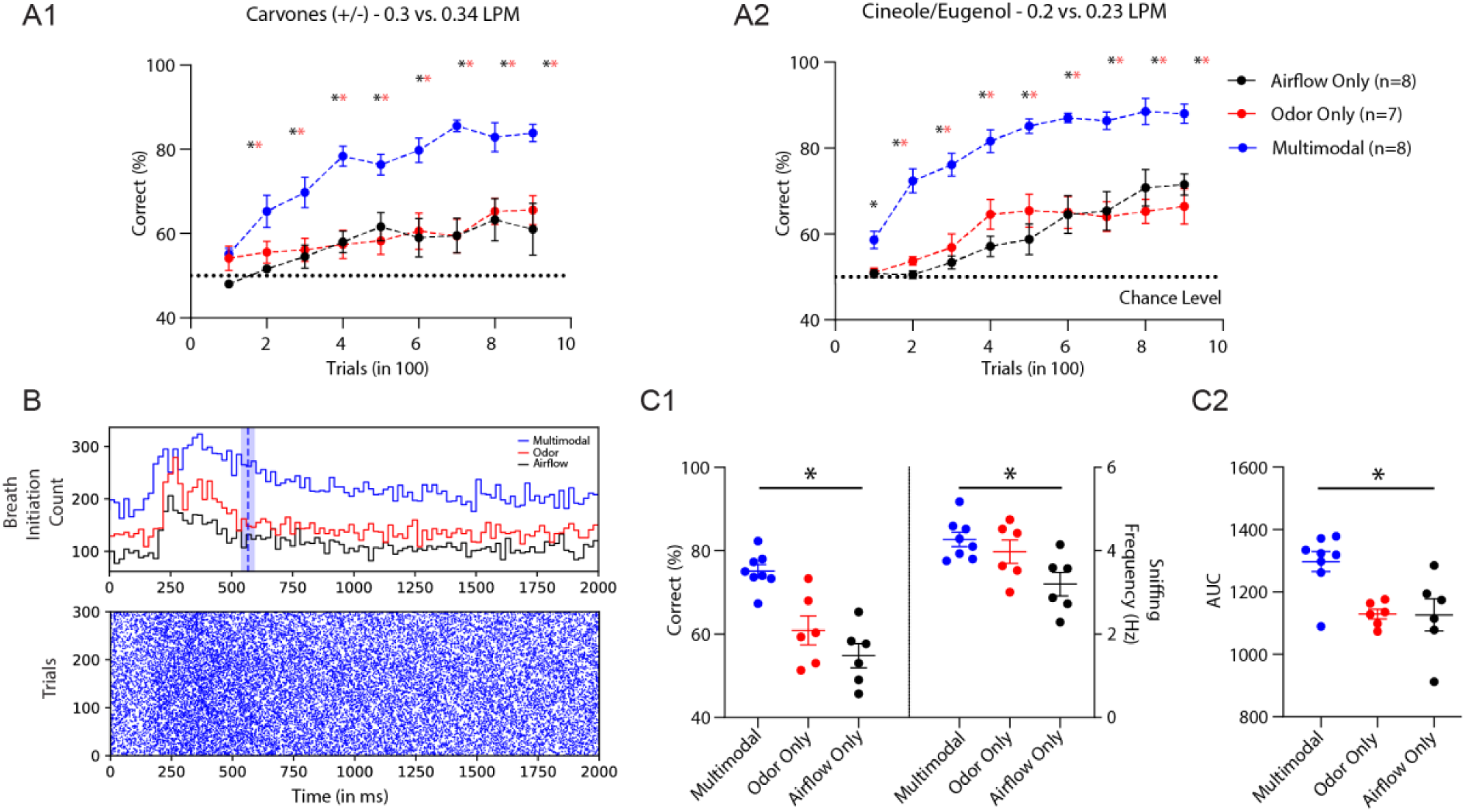
Multimodality generated by airflow-odor combination enhances discrimination learning at the subthreshold levels. **A. A1 and A2.** Comparisons of learning curves of animals when trained to discriminate the unimodal stimuli (odors and airflows) and multimodal stimuli (odor + airflow). For different discrimination tasks animals displayed enhanced learning while they were trained on multimodal stimuli. For Carvones (+/-) - 0.3 vs. 0.34 LPM: Two-way ANOVA, F = 80.96, p<0.0001, Cineole/Eugenol - 0.2 vs. 0.23 LPM: Two-way ANOVA, F = 144.1, p<0.0001, n_Odor_ = 7, n_Airflow_ = 8, n_Multimodal_ = 8). **B.** Raster plots representing inhalation onset for multimodal discrimination task pooled across animals in the last phase of learning (last 300 trials). The histogram representing pooled inhalation onset for odor and airflow only as well as multimodal group. The blue vertical line represents the discrimination time for animals trained on multimodal stimuli and the shaded region represents SEM. **C. C1.** Comparisons of accuracy (Left y-axis) and sniffing frequency (Right y-axis) of animals when trained to discriminate the odor, airflow and multimodal stimuli during the last task. The average accuracy of animals during the last task for multimodal stimuli was significantly higher (One-way ANOVA, F = 17.55, p<0.0001). The average sniffing frequency of animals during the last task for multimodal stimuli was also significantly higher (One-way ANOVA, F = 5.452, p = 0.0148). **C2.** Comparisons of AUC calculated from breathing of animals trained to discriminate the odor, airflow and multimodal stimuli during the last task). The average breathing AUC of animals trained on multimodal stimuli was significantly higher in comparison to those trained on unimodal tasks (One-way ANOVA, F = 7.946, p = 0.0037).

## Discussion

The subsystems present in rodent nose make it a unique sensory organ by imparting the abilities to detect chemical, mechanical and thermal stimuli, yet the neural mechanisms underlying ‘multimodal’ olfaction remain elusive (Ma, 2010). Our experimental results show that mice can discriminate various airflow rates with ∼90% accuracy even in the absence of whiskers, through their nose, proved by olfactory sensory neurons’ (OSNs) ablation, and bulbectomy (Figures 1 and 2). During the anemo-discrimination decision-making, mice exhibited the sniffing refinement indicating the orthonasal airflow information processing (Figure 3). Further, the stimulus-dependent, and learning-dependent calcium signals revealed the role of OB inhibitory circuits in processing airflow information (Figure 4). Genetic/optogenetic modulations resulted in the bidirectional shift of anemo-discrimination learning pace, with contrasting phenotypes for olfactory learning, setting the optimal inhibition for refining odor and airflow stimuli (Figure 5). Furthermore, an enhancement in discrimination learning pace was achieved by combining odor and mechanical stimuli at subthreshold levels, confirming the heightened olfactory perception aided by mechanical stimuli (Figure 6). Thus, revealing the underlying neural mechanisms of mechanosensation through the olfactory system provide a new dimensionality to the olfactory perception.

### Does rodent olfactory system encode and refine airflow information?

Here, we are reporting significant differences between odor and airflow discrimination behavioral readouts. In a go/no-go paradigm employed for investigating odor discriminations, we observed differences in the time taken by animals to discriminate simple and complex odors. The complexities of stimuli were quantified by assessing the overlap between evoked glomerular activity patterns (Abraham et al., 2010; Abraham et al., 2012; Abraham et al., 2004; Abraham et al., 2014; Bhattacharjee et al., 2019). However, when we trained mice to discriminate airflow rates on a similar paradigm, we did not observe any notable differences in FDT across different airflow pairs. We altered the difficulty by varying the discriminable differences and the strength of airflow rates (Figure 1D). Given the fact that reaction time measurements can provide the temporal limit of underlying neural mechanisms, these results may indicate a broader activation pattern evoked in the OB (Wu et al., 2017), independent of stimuli complexity. One commonality we observed with the odor discrimination tasks is the modulation of learning pace and reaction times by the OB inhibitory network, however, strikingly in opposing directions. For example, enhancement of inhibition caused faster learning pace in odor discriminations (Gschwend et al., 2015), whereas it resulted in slow pace for anemo-discriminations (Figure 5F, G). This may arise due to the differences in optimal inhibition required to refine the broader activation patterns evoked by airflows in the OB compared to odorants (Wu et al., 2017). The sniffing modulation observed during the airflow discrimination ratified the orthonasal airflow information processing (Figure 3). The enhancement in sniffing towards the airflow compared to odor stimuli may indicate a strategy used by animals to have optimal activations needed for effective mechanosensation. However, the question of whether OB has the potential to encode and refine the airflow information remained open.

The c-Fos activation observed in GABAergic interneuron population followed by airflow discrimination training, prompted us to analyze the Ca^2+^ dynamics, and we observed learning-dependent refinement in the interneuron activity (Figure 4). As the anemo-information processing by the OB circuits was never reported before, we adopted multipronged experimental strategies to investigate the causality. Deleting GluA2 subunit of AMPARs that causes higher Ca^2+^ influx and enhanced inhibition (Abraham et al., 2010), resulted in slower learning pace, which was supported by the ChR2 stimulation experimental results as well. However, the Arch stimulation resulted in faster learning pace, establishing the causality between inhibitory interneurons and anemo-discrimination (Figure 5). Our study thus provides experimental evidence for the encoding and refinement of airflow information by the rodent olfactory system, an idea that existed for the last 70 years (Adrian, 1951; Ueki and Domino, 1961).

### Mechanosensation at the peripheral olfactory system

As per the reports, approximately 50% OSNs showed mechanical sensitivity, studied under in vitro conditions, where the odor receptors (ORs) have been proposed as the sensors (Connelly et al., 2015; Grosmaitre et al., 2007). Many GPCRs have been shown to be multimodal sensors executing different functions (Caterina et al., 1997; Dhaka et al., 2009; Hilliard et al., 2005). The loss of mechanosensitivity caused either by adenylyl cyclase inhibitor or knocking out the cyclic nucleotide-gated channel CNGA2 indicated the role of second-messenger pathway that is involved in odorant information processing, in mediating mechanosensation (Grosmaitre et al., 2007). Moreover, OSNs that express different ORs showed variable sensitivity towards mechanosensation (Connelly et al., 2015). All these observations point to a plausible sensing mechanism by ORs.

The deformation of neuronal membrane by mechanical stimuli evoked by sniffing modulations and aerodynamics in the nasal cavity, may result in enough conformational changes in mechanosensitive ORs, that ultimately lead to the second messenger cascades. However, in our experiments, we observed a complete shutdown of mice’ ability to discriminate airflows, after ZnSO_4_ and methimazole treatments. If only 50% of OSNs show the mechanical sensitivity, how would a partial OSN ablation (Figure 2C) lead to a complete loss of anemo-discrimination capability? It is possible that additional mechanosensors are present in the olfactory epithelium, which are yet to be discovered. Earlier experiments in the tracheotomized rats proved that the OB neurons fire in accordance with the modulations of airflow through the nasal cavity (equivalent for sniffing) while the animals breathe at a basal rate (Sobel and Tank, 1993). This supports the notion of respiratory modulation acting as a reafferent signaling in the perception. Indeed, the mechanosensation through the OSNs may form the basis for this reafferent signaling.

### Olfactory bulb inhibitory circuits refining the anemo-information processing, a novel function of mouse olfactory system

The enhanced sniffing observed during airflow discrimination could cause an increase in the spiking activity of mechanosensitive neurons. In complex olfactory environments, where animals are challenged to detect and discriminate volatile compounds from odor plumes, this additional channel of information can increase the sensitivity of the olfactory system. This was evident in our experiments, where we observed a significant increase in the learning pace of animals along with heightened sniffing, while mixing two subthreshold stimuli (odor and airflow) (Figure 6). Rhythmic activity has been reported in the OB neurons even in the absence of odor molecules (Onoda and Mori, 1980b). Is this driven by mechanosensitive neurons at the periphery? Or, are there mechanosensors in the OB? In a recent article, the authors report local field oscillations evoked by vascular pressure pulsations. This was mediated by mechanosensitive ion channels present in a subset of OB projection neurons (Jammal Salameh et al., 2024). All these observations lead to the fundamental question, can OB circuits process and refine the mechanical stimuli? If so, how?

Odor encoding by OB projection neurons’ temporal patterns of spiking is very well established (Bathellier et al., 2008b; Cury and Uchida, 2010; Friedrich, 2013; Hopfield, 1995; Laurent, 2002; Margrie and Schaefer, 2003; Schaefer and Margrie, 2007). Further, this activity can be driven by nasal respiration (Carey and Wachowiak, 2011; Cury and Uchida, 2010; Khan et al., 2008). Sniffing modes employed by animals can help them to capture olfactory ‘snapshots’ (Cury and Uchida, 2010; Wachowiak, 2011; Wesson et al., 2008a; Wesson et al., 2008b). The glomerular-specific oscillatory activity recorded from MTCs was proposed to be caused by the mechanosensory channel. The phase of these oscillations remained unchanged across different airflow rates. However, the introduction of odors caused the phase shifts, which were lost in the absence of mechanical stimuli, implying the role of mechanosensation in phase coding of odor information (Iwata et al., 2017). Disruption of inhibition in the OB causes alterations in the phasing of spikes in the respiration cycle, which ultimately lead to impaired odor discrimination (Godde et al., 2016). These observations indicate a probable correlation between OB inhibition and mechanosensory information processing, if OB circuits are involved. Given all these factors, we modulated the synaptic inhibition in different ways. Deleting the GluA2 subunit of AMPARs, or optogenetically activating GAD-65-expressing neurons, the major class of inhibitory interneurons (Parrish-Aungst et al., 2007), which causes enhancement of inhibition, resulted in compromised anemo-discrimination learning. However, inhibiting these GABAergic neurons helped animals to learn better. Moreover, animals showed contrasting learning phenotypes for odor discriminations with these modulations, setting the optimal inhibition needed for the refinement of odor and airflow information (Figure 5). Considering the heterogenous interneuron population in the OB (Burton, 2017; Lledo et al., 2008), and their potential involvement in various disease conditions (Li et al., 2019; Pardasani et al., 2023), it is essential to probe ‘multimodal’ olfaction in smell disorders as well as in the animal models for these conditions (Bhattacharjee et al., 2020; Bhowmik et al., 2023; Croy et al., 2014; Doty, 2007, 2017; Mahajan et al., 2023; Pandey et al., 2024; Pardasani and Abraham, 2022).

Learning-dependent refinement of sniffing behavior was observed during odor discriminations (Bhattacharjee et al., 2019), and anemo-discriminations (Figure 3). Further, a huge increase in the learning pace was observed along with enhanced sniffing, while animals were challenged to discriminate sub-threshold multimodal odor-airflow stimuli compared to unimodal stimuli of odor or airflow (Figure 6). Can these effects be achieved solely by the mechanosensory drive through the OSNs? Or, do we have to probe beyond the sensory periphery? Recent articles (Jammal Salameh et al., 2024; Wang and Hamill, 2021; Zeppilli et al., 2021), along with our analysis using FM1-43 dye uptake, which has been shown to be Piezo2 activity-dependent prove the presence of mechanosensitive ion channels in a subset of OB projection neurons (Supplementary Figure S8) (Coste et al., 2010; Drew and Wood, 2007; Villarino et al., 2023). Given the findings of vascular pressure pulsations driving the spiking activity of mitral cells in the OB through mechanosensitive channels, we propose the anemo-information processing loop, involving the plausible intracranial pressure sensation in the OB through Piezo2 channels, driven by animal’s sniffing behavior. The novel function of olfactory system we report here may help revealing the mechanisms of improved perception and cognition caused by integrative body-mind training where better physiological parameters of heart rate, and breathing are observed (Tang et al., 2009).

## Methods

### Animals Used

A total of 174 normally weaned C57BL/6J males (6-10 weeks at the beginning of the experiments) were utilized for all the experiments (experiment wise breakup of animals is mentioned in the corresponding data). In addition to the wild type animals, transgenic mice were obtained by crossing the following genotypes that were obtained from The Jackson’s laboratory, unless mentioned otherwise.

GluA2^2Lox^: GluA2^2Lox^ (Gria2) was obtained from the Heidelberg University, Germany

GluA2^GAD65(+/-)^ knockout: GluA2^2Lox^ with B6N.Cg-Gad2^tm2(cre)Zjh^/J for the knockout of GluA2 in GAD65+ve neurons.

GAD65-EYFP: B6N.Cg-Gad2^tm2(cre)Zjh^/J with B6.129X1-Gt(ROSA)26Sor^tm1(EYFP)Cos^/J for the expression of EYFP in GAD65+ve cells.

GAD65-GCaMP6f: B6N.Cg-Gad2^tm2(cre)Zjh^/J with B6;129S-Gt(ROSA)26Sor^tm95.1(CAGGCaMP6f)Hze^/J for the expression of GCaMP6f in GAD65+ve cells.

GAD65-Arch: B6N.Cg-Gad2^tm2(cre)Zjh^/J with B6;129S-Gt(ROSA)26Sor^tm35.1(CAGaop3/GFP)Hze^/J for the expression of Archaerhodopsin (Arch) in GAD65+ve cells.

GAD65-ChR2: B6N.Cg-Gad2^tm2(cre)Zjh^/J with B6;129S-Gt(ROSA)26Sor^tm32(CAGCOP4*H134R/EYFP)Hze^/J for the expression of Channelrhodopsin (ChR2) in GAD65+ve cells.

A total of 23 GluA2^2Lox^ (5 for Western blot and 18 for discrimination learning), 22 GluA2^GAD65(+/-)^ Knockouts (6 for Western blot and 16 for discrimination learning), GAD65-EYFP (6 for c-Fos staining and 14 for optogenetics), 12 GAD65-GCaMP6f (calcium imaging), 15 GAD65-Arch (optogenetics) and 10 GAD65-ChR2 (optogenetics) animals were used in different experiments.

### Ethical Approval

The experimental procedures used in this manuscript are approved by the Institutional Animal Ethics Committee (IAEC) at IISER Pune, and the Committee for the Control and Supervision of Experiments on Animals (CCSEA), Government of India (animal facility CCSEA registration number 1496/GO/ReBi/S/11/CCSEA). The usage of animals in this manuscript was approved under protocol numbers IISER/IAEC/2017-02-006 and IISER/IAEC/2017-02-008.

### Maintenance of animals used in the study

All experiments were conducted on adult male mice. The age of mice at the beginning of the experiment was 6-8 weeks. Mice were housed in individually ventilated cages (IVC) in a temperature- and humidity-controlled animal facility on a 12-hour light-dark cycle. The mice were given standard rodent bedding and nesting material. All the experiments were performed during the light cycle. On the days of behavioral training, mice were fed *ad libitum* food, but subjected to a water restriction schedule, however, designed to keep them at >80% of their original body weight. The water restriction schedule was never more than 12 hours long.

### Surgical Procedures

#### Whisker Trimming

Animals were mildly anesthetized with isoflurane and whiskers were trimmed in a row wise fashion using scissors. Whisker trimming was performed once a week on the experimental animals. Animals that were on a water restriction schedule were given free water for two hours before and after the treatment.

#### Whisker Plucking

The process for whisker plucking was identical to that for whisker trimming. Instead of clipping the whiskers, tweezers were used to pluck them from the root.

#### Intranasal Zinc Sulphate injections

Animals were mildly anesthetized with isoflurane and were held with their snout facing upwards. Using a micropipette, 25 µl of 5% zinc sulphate was infused into one of the nostrils. The animal was turned upside down immediately after the infusion to reduce the passage of zinc sulphate into the trachea or lungs. Before infusing the second nostril, the animals were given a two-hour recovery period. After infusing both nostrils, the animals were given a two-day recovery time with *ad libitum* access to food and water. Animals that were on a water restriction schedule were given free access to water for 12 hours before the treatment.

#### Methimazole injection

75 mg/kg of the methimazole dissolved in PBS was injected intraperitoneally (IP injection). Animals were given a two-day recovery period following injections. Animals on a water restriction schedule were given free access to water for 12 hours before the treatment.

#### Surgical aspiration of Olfactory Bulb (Bulbectomy)

The animals were anesthetized with a combination of ketamine and xylazine. The anaesthetized animal was then mounted on the stereotaxic instrument. An incision was performed on the skull over the olfactory bulb area, and the skull was completely cleaned. A 2.5 mm diameter cranial window was formed over the olfactory bulb region using a biopsy punch. A 21 - G needle connected to a vacuum pump was used to aspirate the olfactory bulb. The skin was sutured back together, and the animal was given a two-week recovery time before continuing with the experiments. All of these steps were same for the animals that had sham surgery, except for the excision of the olfactory bulb.

#### Head post implantation for head-restrained behavioral training

The head post was surgically implanted on top of the animal’s skull. First, the animal was anesthetized with an I.P. injection of Ketamine (50 g/g body weight) and Xylazine (10 g/g body weight) and was mounted on the stereotaxic instrument. A local anaesthetic gel (Lignocaine hydrochloride) was applied to the skin over the skull and an incision was made. Using a cotton swab, the region was thoroughly cleaned with artificial cerebrospinal fluid (ACSF) (125 mM NaCl, 5 mM KCl, 10 mM Glucose, 10 mM HEPES, 2 mM CaCl2, 2 mM MgSO4, in 1 L sterile Distilled water, pH = 7.4). The periosteum of the skull was carefully scraped off with a scalpel blade. The skull was allowed to air-dry and an etching agent was applied on the exposed skull for 15 seconds. A liquid primer was then applied to the dried skull using the applicator and the primer was allowed to air-dry for 10 seconds, followed by UV light exposure for 30 seconds. The primer serves as an adhesive for the UV polymerizing cement to adhere to the bone.

A very thin layer of UV cure dental cement was spread on the skull and polymerized using UV light for 20 - 30 seconds, on which the head post was mounted and was exposed to UV for curing. The remaining part of the skull was filled with cold cure acrylic dental cement. The mouse was unmounted from the stereotaxic instrument and placed in the home cage. The animal was given a recovery period of two days.

#### Cranial window and LED implantation for Optogenetics

Blue or amber LEDs were implanted over the OB region to achieve optogenetic modulation of OB specific GAD65-expressing interneurons. To avoid direct contact with brain tissue, the LEDs were implanted over cranial windows. The animals were first anaesthetized with a combination of ketamine and xylazine and was placed on the stereotaxic instrument. An incision was made over the skull, and the skull was thoroughly cleaned. A window 2.5 mm in diameter was created over the olfactory bulb region using a biopsy punch which was then covered by a glass coverslip by sealing its side with cold cure dental cement. On the skull, a head post was implanted posterior to the cranial window. The clarity of the cranial window was checked after a week, and an LED was mounted over it with cold cure dental cement.

#### Implantation of imaging cannula for in-vivo calcium imaging

The animals were anaesthetized with a combination of ketamine and xylazine and 1mm diameter cranial window was created in the centre of the right hemisphere of the OB using a dental drill. Precautionary, a blunt Hamilton needle was lowered 1mm ventral to the dura in the OB and kept as such for 5 minutes to create a path before insertion of the GRIN lens. 1 mm was chosen based on the observation that most of the GAD65 expressing interneurons lies at this depth. The 1 mm protruding GRIN lens was lowered vertically until it reached the GCL. To secure the lens assembly to the skull surface, a combination of cyanoacrylate gum and acrylic dental cement was used. A headpost was implanted behind the implanted lens and animals were given a month to recover.

#### Stereotaxic Injection of FM 1-43 dye

FM 1-43 dye that specifically labels piezo2 activity was injected in the olfactory bulb of the wild type mice using stereotax (Villarino et al., 2023). The animal was first anesthetized with an I.P. injection of Ketamine (50 g/g body weight) and Xylazine (10 g/g body weight) before being mounted on the stereotaxic instrument. A cranial window of approximately 2.5 mm diameter was made above the olfactory bulb. The region was thoroughly cleaned using artificial cerebrospinal fluid. The intersection of the inferior cerebral vein with the hemispheric midline and Bregma was put in the same plane. 12-14 injections were administered in each olfactory bulb at different locations specifically targeting the dorsal and lateral Mitral cell layer in each location. Approximately 100-150 nL of FM1-43 dye (Invitrogen, T3163, diluted in PBS) was injected in each location. The FM 1-43 dye intake was imaged by imaging the fluorescence emission of the dye at ∼580 nm.

### Operant Conditioning Procedures

#### Behavioral training under freely moving conditions

##### Apparatus

Custom modified eight channel freely moving olfactometers (from Knosys, Washington) (Bodyak and Slotnick, 1999) were used for airflow/olfactory discrimination experiments. The instruments were controlled by a customized program written in IGOR (Wavemetrics). The device included an operant chamber where the animals were placed for behavioral training. A circular sampling/reward port secured by an IR beam was located on the right side of the operant chamber. Animals could start a trial by inserting their head and breaking the IR beam that guarded the sampling/reward port. The animal was provided with a stimulus (airflow/odorants) once the trial was initiated. The precise onset of the stimulation was assured by a system of solenoid valves controlled by software. Airflow rates were also controlled by flowmeters connected to the olfactometer channels. Three different modes of stimulus delivery were used based on the nature of the behavioral experiment (Figure 1 and Supplementary Figure S1).

Mode 1 – In this mode, the stimulus was delivered on the snout of animal from the top (Figure 1A). The stimulus was delivered by circular tube of 4 mm diameter. The distance between the delivery tube and the snout was kept fixed at around 8-10 mm. The input airflow rate supplied and the actual output rate observed for mode M1 were quantified and found similar (Supplementary Figure S1A).

Mode 2 – In this mode, the stimulus delivery was targeted on the whiskers. Two protruding tubes (4 mm in diameter) delivered focused stimulation to both sides of the snout (Figure 1E1). The delivery tubes protruded sufficiently to be in the same plane as the lick tube. When the animals poked their heads into the sampling port, they were provided with the airflow stimulus on their whiskers. The radial distance between the lick port and each tube was 8 mm. The delivered input airflow rate and the observed output rate for mode M2 output tubes were quantified and found to be equivalent (Supplementary Figure S1B).

Mode 3 – In this mode also, the stimulus delivery was targeted on the whiskers but in a diffused way. The stimulus was provided to the whiskers by four holes (4 mm diameter each) arranged in a semi arc around the lick tube (Figure 1E2). In this mode, the plane of the lick tube was protruded in comparison to that of the delivery ports. When mice poked their head in the sampling port, they received the stimulus in a diffusive manner. The radial distance from the centre of lick port to that of each hole was also 8 mm in this mode. The real output rate for different output holes of mode M3 was quantified, and the actual observed output was found to be as expected (Supplementary Figure S1C). For Mode M2 and M3, the airflow rates mentioned during the discrimination training are the total output rates from either side i.e. 0.6 LPM means that animals received 0.6 LPM from both sides of the snout.

#### Task Habituation Phase

Three to four days after the start of water deprivation schedule, animals underwent a task habituation training. Animals were trained using standard operant conditioning procedures (Abraham et al., 2004). The task habituation was completed in nine phases. The animals received water reward (2-3 µl) simply by breaking the IR beam in the first pre-training phase (Phase 0). This enabled the animals to locate the reward port and the lick tube. In the next stage the animals only received water when they registered at least a single lick. After 15 such trials, the subsequent phase of task habituation begins wherein the trial is initiated only when IR beam is broken by nose poke. The beam break causes the instrument valves to open, resulting in stimulus presentation for 2 seconds. The task’s complexity was steadily increased from this stage onwards in order to train the mouse to lick continuously during stimulus presentation. Airflow with precise flowrate and diluted odors were delivered for the airflow and odor-based pre-training tasks, respectively. Unless specified otherwise, odors with a concentration of 1% were used and diluted with mineral oil. All animals completed the pre-training in three to four sessions of 30 minutes.

#### Discrimination Training

All the airflow and odor-based discrimination training was performed on a Go/No-Go behavioral paradigm (Abraham et al., 2004; Bodyak and Slotnick, 1999). The mouse initiated a trial by breaking an IR beam that guards the sampling port (Figure 1A). This enabled the opening of one of the solenoid valves, followed by the opening of a three-way diversion valve after 500 milliseconds. After diversion valve is opened, the stimulus is presented to the animal for a specified duration. The use of a diversion valve reduced the period between the onset of the stimulus and the first contact with the animal. To obtain a reward, the animal had to meet the required reward criteria based on the reward contingency of the stimulus [Rewarded (S+)/ Non-Rewarded (S–)].

The response time provided to the animal was virtually divided into four equal bins. (For example, if the given response duration is 2 seconds, it will be virtually divided into four 500 ms bins). The response time was generally kept the same as the stimulus duration. For an S+ trial to be correct animal had to register a lick in at least three of these four bins. For a correct S+ trial animal received a water reward of 3-4 µl after the end of the stimulus [Reward Criterion: Animal needs to register a lick in at least three out of four bins]. To be successful in an S– trial, the animal should not lick in more than two bins. A subsequent trial did not begin unless an Inter trial interval (ITI) of 5 seconds had passed between trials. The set ITI was long enough for the mouse to retract fast at the end of a trial. There were no regulations in place to compel the mouse to sample the stimulus for a minimum time before making a decision. Furthermore, there were no rules in place to restrict licking during the pre-stimulus time.

Stimuli were delivered to the animals in blocks of 20 trials. Each of these blocks contained ten S+ and ten S– trials. The S+ and S– trials within a block were pseudorandomized so that no more than two stimuli of the same reward contingency were delivered consecutively. The preference for a certain stimulus was avoided by counterbalancing the S+ and S– stimuli for a group of animals. Animals were sufficiently motivated to complete 200-300 trials each day, spread out over 1-2 (20-30 min) sessions. Different instrumental readouts, such as inter trial interval and licking frequency, were used to track the motivation of animals. When the animal ceased licking for the rewarded trials, the training session was stopped.

#### Data Acquisition

The data was collected using a custom-written software in IGOR-PRO that was compatible with the MCC-CIO-DIO 48 data acquisition card.

#### Video recording of animals

The videos of the animals performing the airflow discrimination task under freely moving conditions were captured with a Redmi 4A smartphone at a resolution of 30 frames per second.

#### Behavioral Training under head restrained conditions

##### Apparatus

The head restrained apparatus was custom-built to deliver air flow/odor as the stimulus. A lickometer, a 10-channel olfactometer, stimulus delivery nozzle, a respiration/breathing sensor, a lick tube, and a PVC tube were all part of the setup. The lickometer serves as an interface between all of the individual components that records and digitizes lick and breath responses of the animal to various stimuli. The olfactometer was equipped with eleven mass flow controllers and electromagnetic solenoid valves enabling experimenter to attain temporal precision during odor delivery. A splitter put in the olfactometer divided a clean air stream and routed it to various mass flow controllers. Air entered odor bottles via 10 small controllers while air from the main controller was regulated by solenoid valves and used as a dilution air stream. The dilution and odorized air streams were combined at the output through a T-tube before entering the odor delivery nozzle. The stimulus was delivered to the nostril of the restrained animal by the nozzle. A preloading duration was set before the odor presentation to guarantee that the odor plume reached a stable state. After the preloading time, the exhaust was turned off allowing the stimulus to be delivered to the animal. This enabled us to obtain a homogeneous stimulus presentation. The equipment consisted of a PVC tube in which the animal was placed, and the head was restrained by screwing the head post onto a custom-built metallic plate attached to the tube. A lick tube was positioned near the animal’s mouth to record the animal’s lick state during the trial. On finishing the trial and completing the reward criteria the animal received water reward from the lick tube. The sniffing behavior of mice performing odor-based decision-making tasks under head-restrained condition was non-invasively monitored using a thermocouple-based pressure sensor placed near one nostril. The instrument was controlled by a LAB-View application that was custom written. During the head restrained training the ITI was kept constant at 13.2 s as this was the ideal inter trial period we observed while training animals under freely moving conditions.

#### Task Habituation Phase

Animals were subjected to task habituation training three to four days after the start of the water deprivation schedule. This was done to ensure that the animals become acclimated to the instrument and the procedures associated with it. Standard operant conditioning approaches were used to train the animals. This task habituation phase was completed in stages. Regardless of the animals’ responses during the first stage, they got a water reward of 3 µl after the presentation of a 200 ms short tone. This was done for 20 trials, after which the delay between tone and reward was raised to 1 second (20 trials) and then 2 seconds (20 trials). This was done to elicit licking behavior in water-deprived animals. Once the animals had learned to wait for the water, a stimulus was given to them. The stimulation lasted 2 seconds and came before the reward. The stimulus was also delivered to the animal during the next stages of the task habituation phase, and they had to learn to lick solely during the stimulus delivery duration. If animals only licked when a stimulus was present, they received a water reward after meeting the reward criteria (Phase 1 – Minimum 80 ms of licking and the lick is registered in at least 1/4 virtually segregated bins, Phase 2 – 120 ms; 2/4 bins and Phase 2.1 – 240 ms; 3/4 bins). If animals licked prior to the onset of the stimulus, the required licking time to obtain the reward was increased to 200% of the above-mentioned criteria. All of the animals completed the pre-training in 4-5 sessions of 30-40 minutes each. When mice completed the training, they learned not to lick during the baseline, or time before the stimulus onset.

#### Discrimination Training phase

Using a Go/No-Go behavioral paradigm(Abraham et al., 2012; Bhattacharjee et al., 2019), animals were trained to discriminate between two different stimuli, one of which was rewarded (S+) and the other was unrewarded (S–). The discrimination task was performed with a constant ITI of 13.2 seconds. The start of a trial causes one of the valves that controls the stimulus to open. The stimulus was deflected to the exhaust for the first 1 sec using a vacuum pump linked to the glass funnel. This was done to mitigate the time difference between stimulus onset and its encounter with the animal. During this one sec the baseline licking was also assessed. The stimulus was presented for 2 seconds and animals had to respond within this time frame to get a reward. The time required to lick to receive the reward for a S+ trial was determined by the baseline licking. If animals did not lick during the baseline, they were required to lick for a total of 240 ms in three (minimum 80 ms in each bin) of four 500 ms time bins. If the animals licked at the baseline, they had to lick twice as long during the stimulus presentation to get the reward and register it as a correct trial. For an S– trial to be correct, the licking of the animal should not exceed a total of 80 ms, regardless of whether it displayed baseline licking or not. There was no punishment for the incorrect trials. Training was carried out as explained under freely moving conditions. The frequency of licking was used to track the motivation of animals. When the animals were not motivated, they ceased licking for the rewarded trials.

**Table 1:** Training sequence of animals for various tasks under different photoactivation conditions.

| Airflow Pair/Odor Pair | Photoactivation |
| --- | --- |
| 0.5 vs. 0.4 LPM (1200 trials) | OFF |
| 0.15 vs. 0.1 LPM (900 trials) | ON |
| 0.3 vs. 0.25 LPM (600 trials) | OFF |
| 0.45 vs. 0.35 LPM (600 trials) | ON |
| 0.6 vs. 0.6 LPM (300 trials) | OFF |
| Octanol (+) (60%) - (-) (40%) vs. | ON |
| Octanol (-) (60%) - (+) (40%) (600 trials): 0.6 LPM for dilution of both stimuli |  |
| Hexanal (60%) Pentanone - (40%) (0.4 LPM) vs. Pentanone (60%) - Hexanal (40%) (0.3 LPM) (600 trials) | ON |
| Carvone (+) (0.6 vs. 0.4 %) (600 trials): 0.2 LPM for dilution of both stimuli | ON |

#### Measurement of breathing parameters

The breathing dynamics of the animal was continuously recorded during behavioral training using a non-invasive airflow pressure sensor placed near one of the nostrils. The sensor delivered real-time analogue signals to the lickometer linked breathing circuit with a resolution of as good as 4 μS. Using a threshold function, the lickometer breathing circuit converted the raw analogue signals to a digital signal. The thresholding transformed the raw signal in binary values of 0 and 1 with 0-1 transition representing and inhalation and 1-0 transition representing an exhalation. On a trial-by-trial basis, these signals were continuously updated in the results file.

#### Data Acquisition

Data was collected using a custom-written LAB View program that was compatible with a National Instruments (PCI 6703) data acquisition board.

#### Behavioral Readouts

##### Learning curve

We used a measure known as the learning curve to visualize the learning pace and its magnitude. The learning curve measures the change in percentage of correct responses as training progresses. Each point on the learning curve indicates the average accuracy of 100 trails [50 S+ and 50 S–] across all animals.

#### Discrimination time

The timepoint, when the licking response of the animals towards S+ and S– stimuli considerably diverge after the stimulus onset, that time is referred to as the discrimination time (DT). The licking behavior of each mouse was monitored to assess the discrimination time. Animal licking behavior was recorded with high temporal resolution and analyzed in time bins of 20 ms and 2 ms for freely moving and head restrained conditions, respectively. The licking behavior changed as a result of learning and was considerably different between the beginning and end of learning. Early in their learning process, when the animals couldn’t discriminate the difference between S+ and S– stimuli, they licked for both the stimuli. As a result, during the initial training phase, the animal’s lick responses to S+ and S– stimuli were comparable. But as soon as they were able to differentiate between two stimuli, they began to selectively lick for S+ trials and avoid licking for S– trials, which caused a divergence in the lick responses between two stimuli. The discrimination time was determined using a sigmoidal curve that represented the difference between the S+ and S– licking responses. The discrimination time was measured task wise i.e. for 300 trials. The statistical comparison of the lick responses between the S+ (150 trials) and S– (150 trials) trials yielded a p-value curve as a function of time. The discrimination time was considered as the last time point where the p-value was less than 0.05.

#### Intertrial Interval (ITI)

Intertrial interval is the time that animal takes between the two consecutive trials. ITIs were used as the proxy for motivation level for the animals. Under freely moving condition, ITIs were variable and depended upon the animals whereas for head restrained behavior, ITI was kept fixed at 13.2 s. For head restrained training, the motivation is monitored based on the lick behavior of the animals.

#### Breathing analysis

The breathing frequency was measured for each trial throughout the stimulus time as well as before, during, and after the decision-making window. Assuming the discriminating time is x ms different windows were defined as follow: Pre-DT from 0-x to 0, During DT from 0 to 0+x, and Post-DT from 13000-x to 13000. Onset of inhalation time points represented a point in the raster plot. A histogram with a bin resolution of 20 ms was used to depict the inhalation onset distribution across all trials and animals. Time of first inhalation after the onset of stimulus was considered as inhalation onset time. The area under the curve was determined using the trial wise histograms plotted for each animal. The AUC was computed just from the start of the stimulus to the discrimination time/stimulus time. The sniffing peak latency was defined as the time corresponding to the mode of distribution of breath initiations. For analysis, custom built algorithms in Python were employed.

#### Immunohistochemistry

##### Hematoxylin and Eosin staining

The procedure for Hematoxylin and Eosin staining was adopted from our previous study (Mahajan et al., 2023). Briefly, the mice were perfused with 1×PBS and 4% paraformaldehyde (PFA) and the nasal cavity was dissected and kept for a week in a 10% EDTA solution to decalcify the bones surrounding the cavity. Then, the nasal cavity was embedded in the paraffin. Using microtome (RM2235, Leica Biosystems), 5-8 μm thick sections were obtained and were transferred to the poly-L-lysine coated slides and the tissues were stained with Hematoxylin and Eosin as described previously(Mahajan et al., 2023). The DPX mounting media was added to the slides and they were mounted with a cover slip and edges of the cover slip were sealed. A brightfield microscope (BX43, Olympus) was next used to examine the sections and capture the images. For each animal, two or three ROIs per section were imaged depending on the quality of the sections from 8-10 sections per animal. The thickness of the OE was measured as the maximum perpendicular length from the basal membrane and compared across different groups.

#### Pizeo2 staining

The mice were transcardially perused using 1×PBS and 4% paraformaldehyde and the brain was dissected and stored in PFA at 4℃ for 24 hours. The brains were then cryopreserved in 30% sucrose solution for 24 to 48 hours. 50-μm thick coronal sections were taken using Cryotome (ThermoFisher) and were collected in 24 well plate containing 1X PBS. The sections were then washed thrice for 5 minutes each with PBST (0.1% Triton-X). Non-specific sites were blocked using 2 hours long incubation with blocking solution (5% NGS and 1% Triton-X in PBS). After blocking, sections were incubated at 4℃ for 16 hours with rabbit-anti-Piezo2 antibody (NBP1-78624, Novus Biologicals) at 1:500 dilution in 1% NGS and 0.1% Triton-X in PBS. Post incubation in primary antibody, the sections underwent three PBST washes for 15 minutes each. The sections were then incubated with secondary antibody (Anti-Rabbit Igg Alexa 488, 18772-1ML-F, 1:1000 in 1% NGS and 0.1% Triton-X in PBS) for 4 hours at room temperature. Post incubation with secondary antibody, three PBST washes of 15 minutes each was given and then the sections were stained with DAPI (1 μM in PBS). Stained sections were mounted on glass slides using VECTASHIELD (Vector Labs, H-1000) mounting media. The sections were imaged using Leica Sp8 confocal microscope.

#### GAD65 staining

We used GAD65-EYFP animals to visualise the distribution of GAD65 (GAD2) interneurons in the mouse brain. Mice were perfused with 1X PBS and 4% PFA, and the dissected brain was stored at 4 °C for 24 hours in 4% PFA. The brains were then cryopreserved in 30% sucrose solution for 24 hours. 50-μm thick coronal sections were taken using Cryotome and collected in 1X PBS containing 24 well plate. Following sectioning, the sections received a 10-minute 1X PBS wash. After incubating with DAPI (1:500) for 10 minutes, the free-floating sections were washed with 1X PBS for an additional 10 minutes. The sections were placed on glass slides and mounted with VECTASHIELD mounting media. The sections were imaged using Leica Sp8 confocal microscope.

#### **c-** Fos staining of olfactory bulb sections

A group of GAD65-EYFP mice (n = 7) were trained to distinguish between 0.2 vs. 0.1 LPM. Three mice were randomly processed for immunohistochemical analyses of c-Fos once they reached the asymptotic phase of learning. Mice were perfused with 1X PBS and 4% PFA, and the dissected brain was stored at 4 °C for 24 hours in 4% PFA. 50 µm floating coronal sections were taken with a vibratome. These sections were washed three times for 15 minutes each with 1X Tris Buffered Saline (TBS). Blocking was performed for two hours to remove any non-specific binding. The blocking solution contained 0.2% Triton-X (Sigma), 7.5% Normal Goat Serum (NGS) (Abcam, ab7481), and 2.5% Bovine Serum Albumin (BSA) (Sigma Aldrich) in TBS. Following blocking, primary antibody was used: rabbit anti-c-Fos (1:500 dilution). The sections were incubated overnight at 4 °C for 13-15 hours. The following day, three TBS washes (15 minutes each) were given. The secondary antibody was then incubated at room temperature for two hours. The secondary antibodies used was anti-rabbit Alexa Flour 594 (1:500 dilution) (Jackson Immunoresearch, USA, code:111-585-003). Three TBS washes (15 minutes each) were given. Finally, the sections were stained with DAPI (Sigma, 1:500 dilution in 1% NGS) for 10 minutes and were mounted on the slides. The sections were imaged using Leica Sp8 confocal microscope.

#### Western Blotting

We used 8–10 weeks old GluA2-Lox (n = 5) and GAD65-GluA2 knockout (n = 6) animals for western blotting (WB). After being dissected, the olfactory bulbs were stored at −80°C until needed for WB. Whole brain lysates were prepared in RIPA buffer supplemented with complete protease inhibitor. BCA protein assay kit was used to perform the protein estimation. 20 μg sample was loaded in each well of an 10% polyacrylamide gel and SDS–polyacrylamide gel electrophoresis (PAGE) was performed. The protein was then transferred to Immobilon-P PVDF membranes. Blocking was then performed with 5% milk/Tris-buffered saline–Tween 20 for 1 hour at room temperature. Primary anti-GluA2 antibody and anti-GAPDH were used to probe the membranes for 16 hours at 4°C at 1:250 and 1:5000 dilutions, respectively. Peroxidase-conjugated AffiniPure Goat anti-rabbit IgG was used as the secondary antibody. It was used at 1:5000 dilution and incubated for 1 hour at room temperature. Using Clarity ECL Western Blotting Substrate, bound antibody was detected and an ImageQuant LAS 4000 was used to digitally capture the image.

#### Micro-endoscopic Ca^2+^ imaging

Two groups of GAD65-GCaMP6f mice were used to perform Ca^2+^ imaging. The first group consisted of 5 animals that underwent calcium imaging while anaesthetized, while the second group consisted of 5-7 animals that underwent Ca^2+^ imaging while awake.

#### Image acquisition

Calcium imaging was performed on animals while they were anaesthetized and awake under head restrained conditions. GAD65+ve interneurons in the OB GCL were imaged using a snap-in fluorescence microscope (OSFM model L, Doric lenses Inc., Canada) mounted on an implanted GRIN cannula (1 mm length). The cannula had a working distance of 80 µm and a focal range of 50 µm. At a frame rate of 10 Hz, a field of view corresponding to 350 µm X 350 µm that was further binned to 2 × 2 times, was imaged. The CE: YAG fluorescence source of 465 nm output was set used with power ranging from to 400-700 mA.

For imaging under anaesthetized conditions, an IP injection of a ketamine-xylazine cocktail was administered to animals before beginning of each session. Before mounting the animals on the setup, their toe reflexes were tested. Each session consisted of 60-80 trials with an ITI of 13.2 seconds. The animals (first group n = 5 mice) were given different airflows with the stimulus of 100, 200, 300, and 400 mL/min, which were presented to them in a pseudorandomized fashion. Twenty trials per animal were conducted and analysed for each airflow.

Animals were subjected to standard airflow-based discrimination tasks under head restraint conditions for imaging under awake state. Animals (second group n = 5-7) underwent a task habituation phase and were trained to discriminate 0.5 vs. 0.4 LPM (4 tasks - 1200 trials). Calcium traces were recorded during the first 40 trials (20 S+ and 20 S–) of each task (300 trials) at the beginning of each task. An external tone TTL signal (200 ms) from the olfactometer synchronised Ca^2+^ imaging to the start with the trial. The imaging for each trial lasted 10 seconds which included the stimulus duration of 2 seconds. The imaging was done at a resolution of 100 ms. The trial-by-trial imaging data was compiled by comparing the trial sequence generated by the olfactometer result file. **Imaging analysis**

Custom Python scripts were used to analyse the images. The population activity was analysed by selecting the regions-of-interest (ROIs) of 300 µm X 300 µm. If there were any frame drops during image acquisition, such trials were identified and excluded from the analysis. The relative changes in fluorescence for each frame were calculated using F(t)/F0 = [F(t) - F0]/F0, where F0 is the mean activity during the baseline. The F0 was calculated trial-wise. The F(t)/F0 amplitude and area under the curve (AUC) of individual animals were measured trial-wise.

#### Optogenetics Experiments

##### Optogenetic modulation of OB specific GAD65 expressing interneurons

To investigate the effects of inhibition on M/T cells and their effects on airflow detection and discrimination behavior, we ontogenetically modulated the activity of OB specific GAD65 expressing interneurons in a bidirectional manner by expressing ChR2 and Arch under the GAD65 promoter. Three different set of animals were used for optogenetic experiments: First group – GAD65-Arch (n = 12-14), Second group – GAD65-ChR2 (n = 7-10), Third group – GAD65-EYFP (n = 12-14). While mice were performing airflow/odor discrimination, GAD65+ve interneurons were photo-stimulated/photo-inhibited. To activate the interneurons, we used 5 ms pulses of blue light (473 nm specific for ChR2) at a frequency of 40 Hz. To inhibit the interneurons, we used 1000 ms pulses of amber light (595 nm specific for Arch) at 1 Hz frequency. For these experiments, the light stimulations were timed to coincide with the onset of the stimulus.

##### Task habituation phase and LED power standardization

After 1 week of LED implantation the water deprivation schedule of animals began. Animals were subjected to task habituation training three to four days after the start of the water deprivation schedule. The operant conditioning paradigm used for these animals was the same as for the head restrained task habituation phase. As optogenetic experiments were carried out to determine whether modulation of GAD65+ve interneurons affects airflow discrimination behavior, and because the extent of transparency of the cranial window may vary between mice, the optimal power required to modulate the majority of GAD65+ve interneuron activity would also vary. As a result, when the animals completed the task habituation, the strength of the LEDs was standardised using the detection task by varying the power of the LEDs during the task. The LEDs were powered by an LED driver, and the stimulus used to standardise the LED power was a 0.2 LPM airflow stimulus. While standardising the LED intensity, we also ensured that tissue temperature did not go beyond the body temperature (37 °C). Using a laser power and energy metre, we measured the surface temperature of the LED and the wattage for each light power setting.

Animals were given the stimulus along with photodiode (LED) activation. The photoactivation-ON trials were interleaved with no-photoactivation trials (20 light ON trials followed by 10 light-OFF trials). Different light intensities were used, and 20-40 trials were performed for each LED intensity. Lick responses of animals were recorded and compared during stimulus presentation under light ON and OFF conditions. The average lick responses of the animals in GAD65-Arch and GAD65-ChR2 groups decreased as the intensity of LED increased (Figure 5E). For subsequent optogenetic experiments, the LED intensities of the animals were chosen so that there was no difference between the light ON and OFF states. whereas the lick responses of the control EYFP animals did not alter (Arch - 2.93 mW and ChR2 - 6.96 mW). Because GAD65-EFYP was used as a control group and expressed EYFP, which is also expressed in GAD65-ChR2, the intensity of light used for optogenetics training for these animals was the same as for the ChR2 group (6.96 mW). At this light intensity, there was no difference in the lick responses of these animals when compared to those in light OFF conditions (Figure 5E).

##### Discrimination training

Animals were trained for airflow/odor discriminations under head restrained conditions, and the paradigm and reward criteria for this task were the same as those mentioned in the section on behavioral training under head restrained conditions. Whiskers of the animals were kept trimmed throughout the experiment to rule out any possibility of whiskers being involved in airflow discrimination behavior. The whiskers were trimmed once a week. The animals were trained for different airflow and odor pairs. The sequence of training is summarized below.

##### Measurement of airflow in nature

The airflow was measured from the potential rodent habitats that includes burrows, grocery shops, waste disposal sites, etc. with the help of a hot-wire anemometer acquired from PCE instruments. The anemometer was kept inside the burrow or the location of potential habitat and average airflow rate for 100s was measured. The measurements were done from multiple locations and were pooled together to compute the airflow range that rodents encounter in the nature.

##### PID measurements

PID measurements were taken to rule out the presence of any consistent odorant cues within the airflow stimulus. High intensity photons colliding with vaporised molecules of odors, if present, would cause a voltage shift in the photo ionic detector during PID measurements. PID measurements were performed for various airflows using a PID probe under similar conditions that was used for airflow detection and discrimination experiments. PID measurements were also performed on a set of odorants in order to demonstrate the difference that exists between airflow and odor generated PID patterns.

##### Data and Statistical analysis

GraphPad Prism 9, Microsoft Excel, and Python were used for all statistical analyses in this study. The data is presented as Mean ± SEM. To determine p-values and test for statistical significance, we used the student’s t-test, one-way and two-way ANOVA, and associated post-hoc tests.

## Acknowledgements

We thank Laboratory of Neural Circuits and Behavior (LNCB) members and IISER-Pune Biology colleagues for fruitful discussions. We thank Profs. Thomas Kuner and Andreas Schaefer for their feedback on the manuscript. We thank staff of *National Facility for Gene Function in Health and Disease* (NFGFHD) and IISER Biology-Leica microscopy facility for the technical support. Some of the illustrations were created with BioRender.com.

## Funding

This work was supported by the DBT/Wellcome Trust India Alliance intermediate grant (IA/I/14/1/501306 to N.M.A.), DBT/Wellcome Trust India Alliance senior grant (IA/S/22/2/506517 to N.M.A.), DST-Cognitive Science Research Initiative (DST/CSRI/2017/271 to N.M.A.), CSIR Fellowship (S.M., S.D., S.D.M., S.P.), UGC NET Fellowship (A.S.B.), and IISER-Pune Fellowship (M.P.). Part of the work was carried at the National Facility for Gene Function in Health and Disease (NFGFHD) at IISER Pune, supported by a grant from the Department of Biotechnology, Govt. of India (BT/INF/22/SP17358/2016).

## Author Contributions

N.M.A. supervised all aspects of the project. N.M.A. carried out the study conceptualization and experimental design. S.M. performed behavioral, optogenetic, Ca^2+^ imaging, and confocal imaging experiments and analysed the data. S.T., S.D., P.S., S.D.M, A.A., L.R., and S.P. helped with various behavioral experiments, immunostaining, stereotaxic surgeries, microscopy and analysis. A.S.B. and M.P. helped with microendoscopic surgeries, Ca^2+^ imaging, and analysis. N.M.A. and S.M. wrote the manuscript with comments from others.

## Competing Interests

The authors have stated explicitly that there are no conflicts of interests in connection with this article.

## Data and Materials Availability

All cumulative data are available in the article/supplementary materials, further inquiries can be directed to the corresponding author.

## List of supplementary material

Supplementary Figures S1 to S8

## Notes

### Competing Interest Statement

The authors have declared no competing interest.

## References

1. Abraham, N.M., Egger, V., Shimshek, D.R., Renden, R., Fukunaga, I., Sprengel, R., Seeburg, P.H., Klugmann, M., Margrie, T.W., Schaefer, A.T., et al. (2010). Synaptic Inhibition in the Olfactory Bulb Accelerates Odor Discrimination in Mice. Neuron 65, 399–411.

2. Abraham, N.M., Guerin, D., Bhaukaurally, K., and Carleton, A. (2012). Similar odor discrimination behavior in head-restrained and freely moving mice. Plos One 7, e51789.

3. Abraham, N.M., Spors, H., Carleton, A., Margrie, T.W., Kuner, T., and Schaefer, A.T. (2004). Maintaining accuracy at the expense of speed: stimulus similarity defines odor discrimination time in mice. Neuron 44, 865–876.

4. Abraham, N.M., Vincis, R., Lagier, S., Rodriguez, I., and Carleton, A. (2014). Long term functional plasticity of sensory inputs mediated by olfactory learning. Elife 3, e02109.

5. Adrian, E.D. (1942). Olfactory reactions in the brain of the hedgehog. The Journal of physiology 100, 459–473.

6. Adrian, E.D. (1951). The role of air movement in olfactory stimulation. J Physiol 114, 4–5p.

7. Alberts, J.R., and Galef, B.G., Jr. (1971). Acute anosmia in the rat: a behavioral test of a peripherally-induced olfactory deficit. Physiol Behav 6, 619–621.

8. Bathellier, B., Buhl, D.L., Accolla, R., and Carleton, A. (2008a). Dynamic ensemble odor coding in the mammalian olfactory bulb: Sensory information at different timescales. Neuron 57, 586–598.

9. Bathellier, B., Buhl, D.L., Accolla, R., and Carleton, A. (2008b). Dynamic ensemble odor coding in the mammalian olfactory bulb: sensory information at different timescales. Neuron 57, 586–598.

10. Bergman, U., and Brittebo, E.B. (1999). Methimazole toxicity in rodents: covalent binding in the olfactory mucosa and detection of glial fibrillary acidic protein in the olfactory bulb. Toxicol Appl Pharmacol 155, 190–200.

11. Bhattacharjee, A.S., Joshi, S.V., Naik, S., Sangle, S., and Abraham, N.M. (2020). Quantitative assessment of olfactory dysfunction accurately detects asymptomatic COVID-19 carriers. Eclinicalmedicine 28, 100575.

12. Bhattacharjee, A.S., Konakamchi, S., Turaev, D., Vincis, R., Nunes, D., Dingankar, A.A., Spors, H., Carleton, A., Kuner, T., and Abraham, N.M. (2019). Similarity and Strength of Glomerular Odor Representations Define a Neural Metric of Sniff-Invariant Discrimination Time. Cell Rep 28, 2966-+.

13. Bhowmik, R., Pardasani, M., Mahajan, S., Magar, R., Joshi, S.V., Nair, G.A., Bhattacharjee, A.S., and Abraham, N.M. (2023). Persistent olfactory learning deficits during and post-COVID-19 infection. Curr Res Neurobiol 4, 100081.

14. Bodyak, N., and Slotnick, B. (1999). Performance of mice in an automated olfactometer: odor detection, discrimination and odor memory. Chem Senses 24, 637–645.

15. Buck, L., and Axel, R. (1991). A Novel Multigene Family May Encode Odorant Receptors - a Molecular-Basis for Odor Recognition. Cell 65, 175–187.

16. Burton, S.D. (2017). Inhibitory circuits of the mammalian main olfactory bulb. J Neurophysiol 118, 2034–2051.

17. Bushdid, C., Magnasco, M.O., Vosshall, L.B., and Keller, A. (2014). Humans Can Discriminate More than 1 Trillion Olfactory Stimuli. Science 343, 1370–1372.

18. Carey, R.M., and Wachowiak, M. (2011). Effect of sniffing on the temporal structure of mitral/tufted cell output from the olfactory bulb. J Neurosci 31, 10615–10626.

19. Caterina, M.J., Schumacher, M.A., Tominaga, M., Rosen, T.A., Levine, J.D., and Julius, D. (1997). The capsaicin receptor: a heat-activated ion channel in the pain pathway. Nature 389, 816–824.

20. Connelly, T., Yu, Y.Q., Grosmaitre, X., Wang, J., Santarelli, L.C., Savigner, A., Qiao, X., Wang, Z.S., Storm, D.R., and Ma, M.H. (2015). G protein-coupled odorant receptors underlie mechanosensitivity in mammalian olfactory sensory neurons. P Natl Acad Sci USA 112, 590–595.

21. Coste, B., Mathur, J., Schmidt, M., Earley, T.J., Ranade, S., Petrus, M.J., Dubin, A.E., and Patapoutian, A. (2010). Piezo1 and Piezo2 Are Essential Components of Distinct Mechanically Activated Cation Channels. Science 330, 55–60.

22. Croy, I., Nordin, S., and Hummel, T. (2014). Olfactory disorders and quality of life--an updated review. Chem Senses 39, 185–194.

23. Cury, K.M., and Uchida, N. (2010). Robust odor coding via inhalation-coupled transient activity in the mammalian olfactory bulb. Neuron 68, 570–585.

24. Dhaka, A., Uzzell, V., Dubin, A.E., Mathur, J., Petrus, M., Bandell, M., and Patapoutian, A. (2009). TRPV1 is activated by both acidic and basic pH. J Neurosci 29, 153–158.

25. Doty, R.L. (2007). Olfaction in Parkinson’s disease. Parkinsonism Relat Disord 13 *Suppl 3*, S225–228.

26. Doty, R.L. (2017). Olfactory dysfunction in neurodegenerative diseases: is there a common pathological substrate? Lancet Neurol 16, 478–488.

27. Drew, L.J., and Wood, J.N. (2007). FM1-43 is a permeant blocker of mechanosensitive ion channels in sensory neurons and inhibits behavioural responses to mechanical stimuli. Mol Pain 3, 1.

28. Fletcher, M.L., and Chen, W.R. (2010). Neural correlates of olfactory learning: Critical role of centrifugal neuromodulation. Learn Memory 17, 561–570.

29. Friedrich, R.W. (2013). Neuronal computations in the olfactory system of zebrafish. Annu Rev Neurosci 36, 383–402.

30. Gerkin, R.C., and Castro, J.B. (2015). The number of olfactory stimuli that humans can discriminate is still unknown. Elife 4.

31. Gire, D.H., and Schoppa, N.E. (2009). Control of On/Off Glomerular Signaling by a Local GABAergic Microcircuit in the Olfactory Bulb. J Neurosci 29, 13454–13464.

32. Godde, K., Gschwend, O., Puchkov, D., Pfeffer, C.K., Carleton, A., and Jentsch, T.J. (2016). Disruption of Kcc2-dependent inhibition of olfactory bulb output neurons suggests its importance in odour discrimination. Nat Commun 7, 12043.

33. Grosmaitre, X., Santarelli, L.C., Tan, J., Luo, M.M., and Ma, M.H. (2007). Dual functions of mammalian olfactory sensory neurons as odor detectors and mechanical sensors. Nat Neurosci 10, 348–354.

34. Gschwend, O., Abraham, N.M., Lagier, S., Begnaud, F., Rodriguez, I., and Carleton, A. (2015). Neuronal pattern separation in the olfactory bulb improves odor discrimination learning. Nat Neurosci 18, 1474-+.

35. Gschwend, O., Beroud, J., and Carleton, A. (2012). Encoding Odorant Identity by Spiking Packets of Rate-Invariant Neurons in Awake Mice. Plos One 7.

36. Hilliard, M.A., Apicella, A.J., Kerr, R., Suzuki, H., Bazzicalupo, P., and Schafer, W.R. (2005). In vivo imaging of C. elegans ASH neurons: cellular response and adaptation to chemical repellents. EMBO J 24, 63–72.

37. Hopfield, J.J. (1995). Pattern recognition computation using action potential timing for stimulus representation. Nature 376, 33–36.

38. Iwata, R., Kiyonari, H., and Imai, T. (2017). Mechanosensory-Based Phase Coding of Odor Identity in the Olfactory Bulb. Neuron 96, 1139-+.

39. Jammal Salameh, L., Bitzenhofer, S.H., Hanganu-Opatz, I.L., Dutschmann, M., and Egger, V. (2024). Blood pressure pulsations modulate central neuronal activity via mechanosensitive ion channels. Science 383, eadk8511.

40. Khan, A.G., Thattai, M., and Bhalla, U.S. (2008). Odor representations in the rat olfactory bulb change smoothly with morphing stimuli. Neuron 57, 571–585.

41. Kleinfeld, D., Deschenes, M., Wang, F., and Moore, J.D. (2014). More than a rhythm of life: breathing as a binder of orofacial sensation. Nat Neurosci 17, 647–651.

42. Laurent, G. (2002). Olfactory network dynamics and the coding of multidimensional signals. Nat Rev Neurosci 3, 884–895.

43. Li, W., Li, S., Shen, L., Wang, J., Wu, X., Li, J., Tu, C., Ye, X., and Ling, S. (2019). Impairment of Dendrodendritic Inhibition in the Olfactory Bulb of APP/PS1 Mice. Front Aging Neurosci 11, 2.

44. Lledo, P.M., Merkle, F.T., and Alvarez-Buylla, A. (2008). Origin and function of olfactory bulb interneuron diversity. Trends Neurosci 31, 392–400.

45. Ma, M. (2010). Multiple Olfactory Subsystems Convey Various Sensory Signals. In The Neurobiology of Olfaction, A. Menini, ed. (Boca Raton (FL)).

46. Macrides, F., and Chorover, S.L. (1972). Olfactory bulb units: activity correlated with inhalation cycles and odor quality. Science (New York, N Y) 175, 84–87.

47. Mahajan, S., Sen, D., Sunil, A., Srikanth, P., Marathe, S.D., Shaw, K., Sahare, M., Galande, S., and Abraham, N.M. (2023). Knockout of angiotensin converting enzyme-2 receptor leads to morphological aberrations in rodent olfactory centers and dysfunctions associated with sense of smell. Front Neurosci 17, 1180868.

48. Mainland, J., and Sobel, N. (2006). The sniff is part of the olfactory percept. Chem Senses 31, 181–196.

49. Margrie, T.W., and Schaefer, A.T. (2003). Theta oscillation coupled spike latencies yield computational vigour in a mammalian sensory system. J Physiol 546, 363–374.

50. Meister, M. (2015). On the dimensionality of odor space. Elife 4.

51. Miura, K., Mainen, Z.F., and Uchida, N. (2012). Odor representations in olfactory cortex: distributed rate coding and decorrelated population activity. Neuron 74, 1087–1098.

52. Mombaerts, P., Wang, F., Dulac, C., Chao, S.K., Nemes, A., Mendelsohn, M., Edmondson, J., and Axel, R. (1996). Visualizing an olfactory sensory map. Cell 87, 675–686.

53. Onoda, N., and Mori, K. (1980a). Depth distribution of temporal firing patterns in olfactory bulb related to air-intake cycles. J Neurophysiol 44, 29–39.

54. Onoda, N., and Mori, K. (1980b). Depth distribution of temporal firing patterns in olfactory bulb related to air-intake cycles. J Neurophysiol 44, 29–39.

55. Pandey, S., Bapat, V., Abraham, J.N., and Abraham, N.M. (2024). Long COVID: From olfactory dysfunctions to viral Parkinsonism. World J Otorhinolaryngol Head Neck Surg 10, 137–147.

56. Pardasani, M., and Abraham, N.M. (2022). Neurotropic SARS-CoV-2: Causalities and Realities. In COVID-19 Pandemic, Mental Health and Neuroscience, P. Sara, and O. Berend, eds. (Rijeka: IntechOpen), p. Ch. 1.

57. Pardasani, M., Ramakrishnan, A.M., Mahajan, S., Kantroo, M., McGowan, E., Das, S., Srikanth, P., Pandey, S., and Abraham, N.M. (2023). Perceptual learning deficits mediated by somatostatin releasing inhibitory interneurons of olfactory bulb in an early life stress mouse model. Mol Psychiatry.

58. Pardo-Vazquez, J.L., Castineiras-de Saa, J.R., Valente, M., Damiao, I., Costa, T., Vicente, M.I., Mendonca, A.G., Mainen, Z.F., and Renart, A. (2019). The mechanistic foundation of Weber’s law. Nat Neurosci 22, 1493–1502.

59. Parrish-Aungst, S., Shipley, M.T., Erdelyi, F., Szabo, G., and Puche, A.C. (2007). Quantitative analysis of neuronal diversity in the mouse olfactory bulb. J Comp Neurol 501, 825–836.

60. Schaefer, A.T., and Margrie, T.W. (2007). Spatiotemporal representations in the olfactory system. Trends Neurosci 30, 92–100.

61. Smear, M., Shusterman, R., O’Connor, R., Bozza, T., and Rinberg, D. (2011). Perception of sniff phase in mouse olfaction. Nature 479, 397–400.

62. Smith, J.C., Ellenberger, H.H., Ballanyi, K., Richter, D.W., and Feldman, J.L. (1991). Pre-Botzinger complex: a brainstem region that may generate respiratory rhythm in mammals. Science 254, 726–729.

63. Sobel, E.C., and Tank, D.W. (1993). Timing of odor stimulation does not alter patterning of olfactory bulb unit activity in freely breathing rats. J Neurophysiol 69, 1331–1337.

64. Tang, Y.Y., Ma, Y., Fan, Y., Feng, H., Wang, J., Feng, S., Lu, Q., Hu, B., Lin, Y., Li, J., et al. (2009). Central and autonomic nervous system interaction is altered by short-term meditation. Proc Natl Acad Sci U S A 106, 8865–8870.

65. Uchida, N., Kepecs, A., and Mainen, Z.F. (2006). Seeing at a glance, smelling in a whiff: rapid forms of perceptual decision making. Nat Rev Neurosci 7, 485–491.

66. Ueki, S., and Domino, E.F. (1961). Some evidence for a mechanical receptor in olfactory function. J Neurophysiol 24, 12–25.

67. Villarino, N.W., Hamed, Y.M.F., Ghosh, B., Dubin, A.E., Lewis, A.H., Odem, M.A., Loud, M.C., Wang, Y., Servin-Vences, M.R., Patapoutian, A., et al. (2023). Labeling PIEZO2 activity in the peripheral nervous system. Neuron 111, 2488–2501 e2488.

68. Wachowiak, M. (2011). All in a Sniff: Olfaction as a Model for Active Sensing. Neuron 71, 962–973.

69. Wang, J.G., and Hamill, O.P. (2021). Piezo2-peripheral baroreceptor channel expressed in select neurons of the mouse brain: a putative mechanism for synchronizing neural networks by transducing intracranial pressure pulses. J Integr Neurosci 20, 825–837.

70. Weber, E.H. (1834). De Pulsu, resorptione, auditu et tactu: Annotationes anatomicae et physiologicae (C.F. Koehler).

71. Wesson, D.W. (2013). Sniffing Behavior Communicates Social Hierarchy. Curr Biol 23, 575–580.

72. Wesson, D.W., Carey, R.M., Verhagen, J.V., and Wachowiak, M. (2008a). Rapid encoding and perception of novel odors in the rat. Plos Biol 6, 717–729.

73. Wesson, D.W., Donahou, T.N., Johnson, M.O., and Wachowiak, M. (2008b). Sniffing behavior of mice during performance in odor-guided tasks. Chem Senses 33, 581–596.

74. Wu, R.Q., Liu, Y., Wang, L., Li, B., and Xu, F.Q. (2017). Activity Patterns Elicited by Airflow in the Olfactory Bulb and Their Possible Functions. J Neurosci 37, 10700–10711.

75. Zeppilli, S., Ackels, T., Attey, R., Klimpert, N., Ritola, K.D., Boeing, S., Crombach, A., Schaefer, A.T., and Fleischmann, A. (2021). Molecular characterization of projection neuron subtypes in the mouse olfactory bulb. Elife 10.

