## Supplementary Material for "Mouse olfactory system acts as anemo-detector and -discriminator"

**The PDF file includes:**
Supplementary Figures S1 to S8

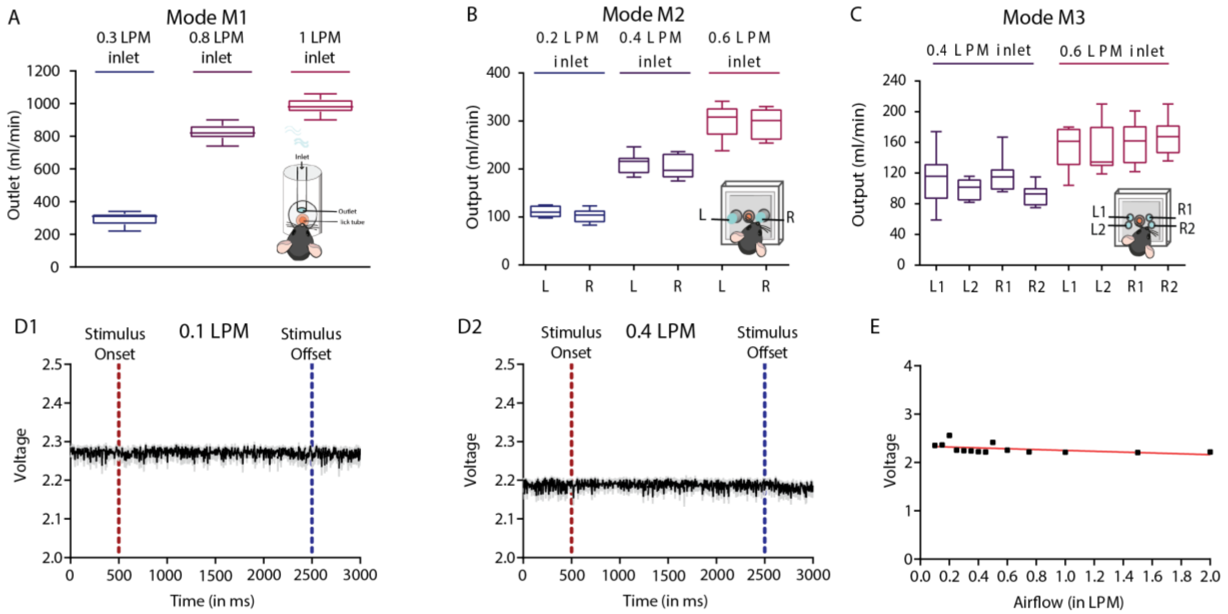


**Figure S1.** Flowrate and PID measurements

A) Outlet airflow was measured from the stimulus delivery tube in the Mode M1 while different airflows were provided as the input. For 0.3 LPM (or 300 ml/min) inlet, the measured output was 296 ± 12.58 ml/min (Mean ± SEM). For 0.8 LPM inlet (or 800 ml/min), output was 793 ± 10.44 ml/min and for 1.0 LPM (or 1000 ml/min) the output was 980 ± 15.78 ml/min. The input airflow rate supplied and the actual output rate observed were similar.

B) Outlet airflow was measured from different ports of the stimulus delivery tube in the Mode M2 while different airflows were provided as the input. For 0.2 LPM (or 200 ml/min) total inlet, the measured output was (Left port – 111.5 ± 3.625 ml/min and Right port – 102.8 ± 4.735), for 0.4 LPM (or 400 ml/min) total inlet, output was (Left port – 212.6 ± 7.131 ml/min and Right port – 203.5 ± 8.360), and for 0.6 LPM (or 600 ml/min) the output was (Left port – 303.3 ± 12.09 ml/min and Right port – 296.5 ± 10.62). The airflow rates through left and right port for a particular inlet stimulus were non-significant and their sum were similar to that of the inlet airflow.

C) Outlet airflow was measured from different ports of the stimulus delivery tube in the Mode M3 while different airflows were provided as the input. For 0.4 LPM (or 400 ml/min) total inlet, the measured output was (Left port1 – 112.6 ± 12.23 ml/min, Left port 2 – 99.25 ± 4.758, Right port 1 – 117.5 ± 8.082 and Right port2 – 92.50 ± 4.679), for 0.6 LPM (or 600 ml/min) total inlet, output was (Left port1 – 154.5 ± 9.716 ml/min, Left port 2 – 151.3 ± 11.40, Right port 1 – 159.6 ± 9.365 and Right port2 – 166.6 ± 8.294). The airflow rates through all the ports for a particular inlet stimulus were non-significant and their sum were similar to that of the inlet airflow.

D) **D1** and **D2**. PID profiles of 0.1 LPM and 0.4 LPM. For both the airflows the voltage in the PID profiles remains unchanged after the stimulus onset.

E) Curve representing change in amplitude voltage of different airflows with respect to their stimulus strength. The amplitude and stimulus strength exhibited no correlation (Pearson $R^2$ = .2176) as well as no linear trend (Linear Regression $R^2$ = .1572).

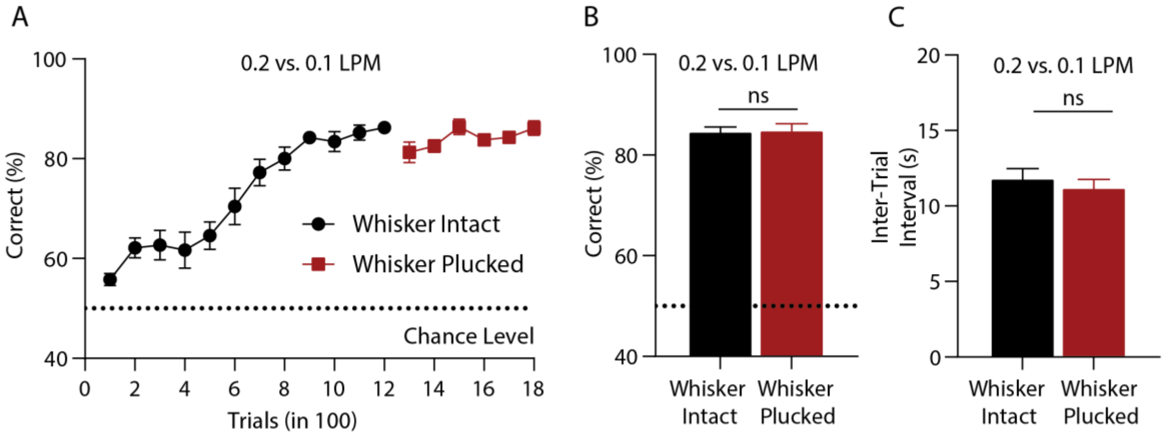


**Figure S2.** Performance of animals on 0.2 vs. 0.1 LPM before and after whisker plucking.

**A.** Learning curve before and after whisker plucking. Animals reached asymptotic phase of learning under both the conditions.

**B.** Final accuracies of animals (average of last 300 trials) before and after whisker plucking. The accuracies were found to be similar (Before: 84.96 ± 1.261; After: 84.71 ± 0.8578, two-tailed paired t-test, p = 0.7550, n = 9).

**C.** Final ITIs of animals (average of last 300 trials) before and after whisker plucking. The ITIs was found to be similar (Before: 11.22 ± 0.5317 s, After: 11.84 ± 0.6322, two-tailed paired t-test, p = 0.3043, n = 9).

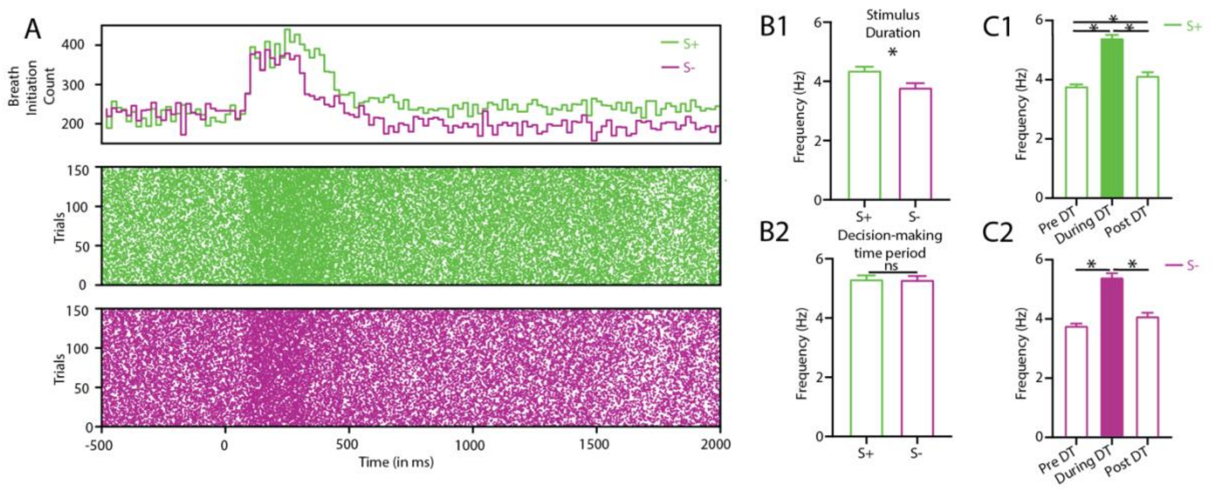


**Figure S3.** Sampling behaviour of animals for S+ and S– trials pooled across airflow pairs

- **A.** Raster plots and histogram representing inhalation onset for S+ and S– trials pooled across all airflow pairs.
- **B.** **B1.** Bar graphs representing sniff frequencies of animals for S+ and S– trials during 2s stimulus duration. The SFs of animals for S+ trials were significantly higher than S– trials (two-tailed paired t-test, $p < 0.0001$, n = 5-8 mice). **B2.** Bar graphs representing sniff frequencies of animals for S+ and S– trials during decision-making window. The SFs of animals for S+ and S– trials were similar (two-tailed paired t-test, $p = 0.8663$, n = 5-8 mice).
- **C.** **C1.** Bar graphs representing sniff frequencies of animals for S+ trials for pre, during and post decision-making period. The SFs of animals during the decision-making period were higher (one-way repeated measures ANOVA with Tukey's multiple comparison test, $F = 84.92$, $p < 0.0001$, n = 5-8). **C2.** Bar graphs representing sniff frequencies of animals for S– trials for pre, during and post decision-making period. The SFs of animals during the decision-making period were higher (one-way repeated measures ANOVA with Tukey's multiple comparison test, $F = 68.19$, $p < 0.0001$, n = 5-8).

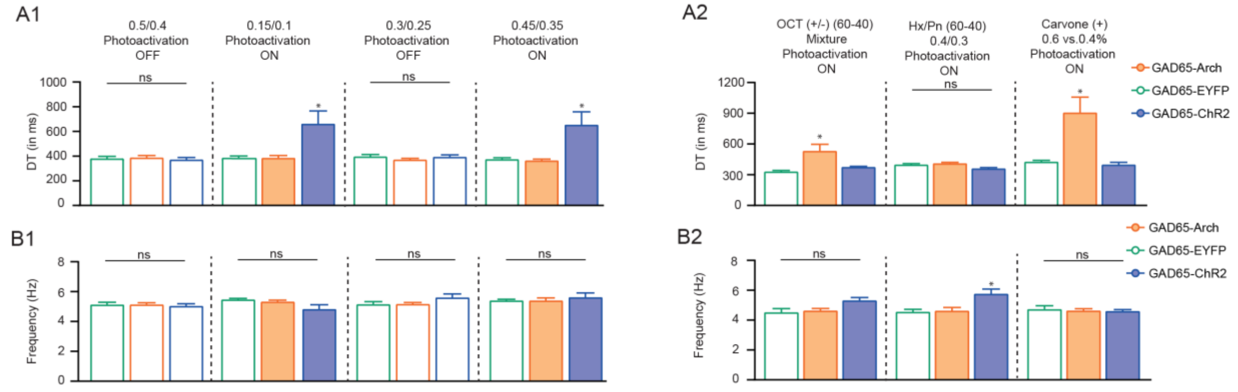

**Figure S4.** Discrimination time and sniff frequency of animals for airflow and odor-based tasks under different photoactivation conditions.

**A.** **A1.** Bar graphs representing average DTs for all groups of animals for various airflows under different photoactivation conditions. The DTs of animals under photoactivation conditions for ChR2 group was slower compared to other groups of animals. (One-way ANOVA with Tukey's multiple comparison test * represents $p < 0.05$). **A2** Bar graphs representing average DTs for all groups of animals for various odour-based tasks. One-way ANOVA with Tukey's multiple comparisons. (* represents $p < 0.05$).

**B.** **B1.** Bar graphs representing average sniff frequencies for all groups of animals during discrimination time for various airflows under different photoactivation conditions. No differences in the SF for all the groups under different photoactivation conditions were observed. (One-way ANOVA with Tukey's multiple comparison test * represents $p < 0.05$). **B2.** Bar graphs representing average sniff frequencies during discrimination time for all groups of animals for various odour-based tasks. One-way ANOVA with Tukey's multiple comparisons. (* represents $p < 0.05$).

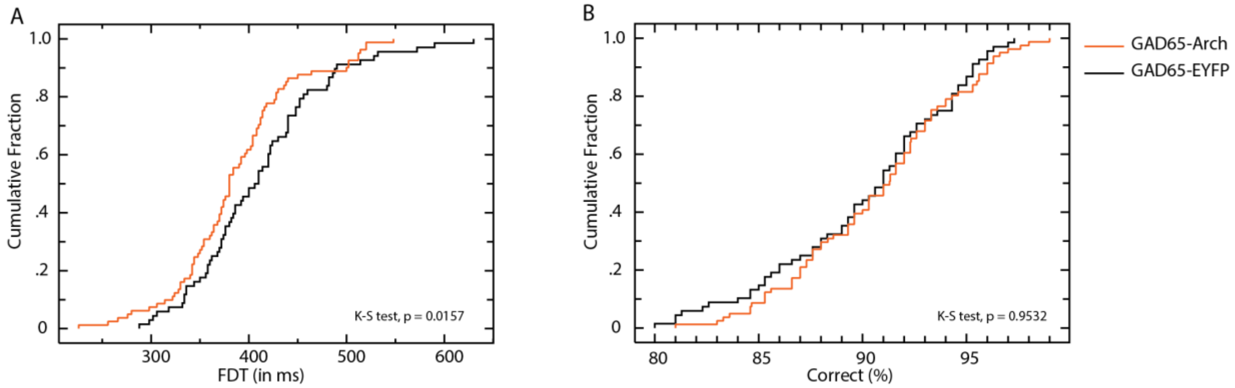


**Figure S5.** GAD65-Arch animals showed faster flow discrimination time (FDT) under photoactivation conditions for airflow discrimination task.

**A.** Cumulative fraction representing the frequency corresponding to a specific DT for different groups. Orange line represents cumulative fraction of DTs for GAD65-Arch group, whereas black line represents the same for GAD65-EYFP group. Under photoactivation conditions Arch animals made faster decisions. (K-S, D-value = 0.256, $p = 0.0157$).

**B.** Cumulative fraction representing the frequency corresponding to a specific accuracy for different groups. Orange line represents cumulative fraction of accuracies for GAD65-Arch group, whereas black line represents the same for GAD65-EYFP group. The accuracy of both groups of animals were similar. (K-S, D-value = 0.0848, $p = 0.9532$).

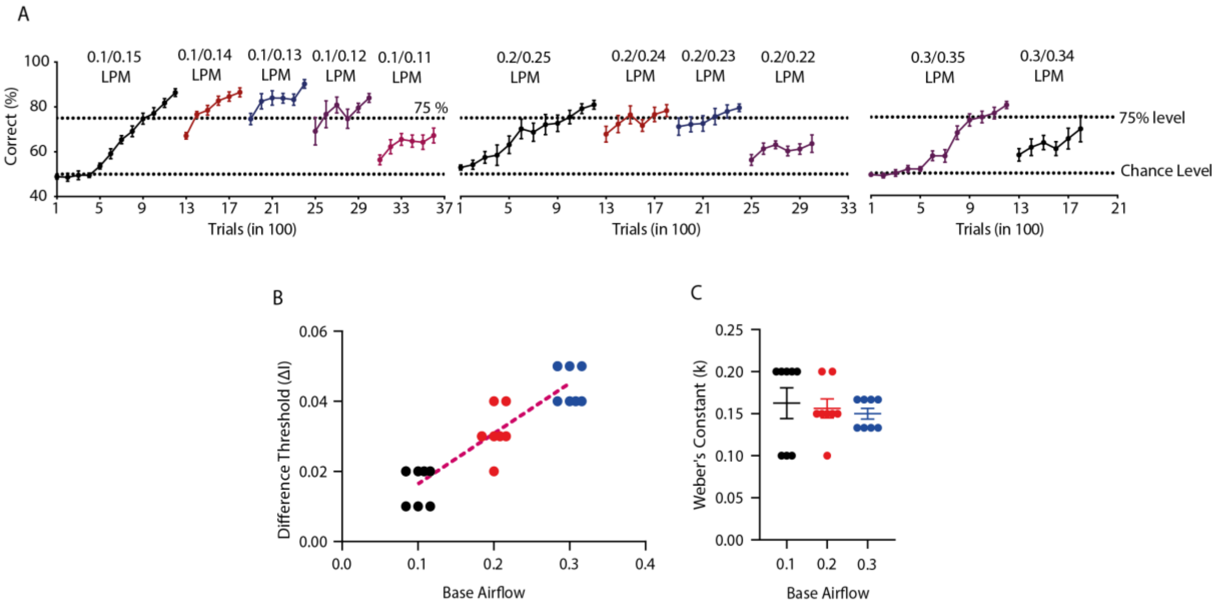


**Figure S6.** Airflow discrimination via olfactory system follows Weber's Law

**A.** Finding difference threshold during airflow discrimination. One groups of animals were trained to discriminate different airflow rates (0.1 vs. 0.15 LPM, 0.1 vs 0.14 LPM, 0.1 vs. 0.13 LPM, 0.1 vs. 0.12 LPM, 0.1 vs. 0.11 LPM) in order to calculate the discrimination threshold or just noticeable difference ($JND_{75}$) for 0.1 LPM, while the difference threshold was kept at 75% accuracy. Similar is done with other two groups of animals that were trained to find the JND for 0.2 and 0.3 LPM as well.

**B.** Difference threshold (ΔI) for different base airflows. Points represent values of ΔI for individual animals. ΔI increases with increase in the airflow base value ($R^2 = 0.9944$, $p = 0.060$). Magenta dotted line shows linear regression with $R^2 = 0.8300$ and slope = 0.1438) (n = 8 for all three base airflows).

**C.** Weber's constant (k) calculated for different base airflows. The dots represent individual values whereas horizontal line represents mean along with SEM (one-way ANOVA, F = 0.2330, p = 0.7941) (n = 8 for all three base airflows).

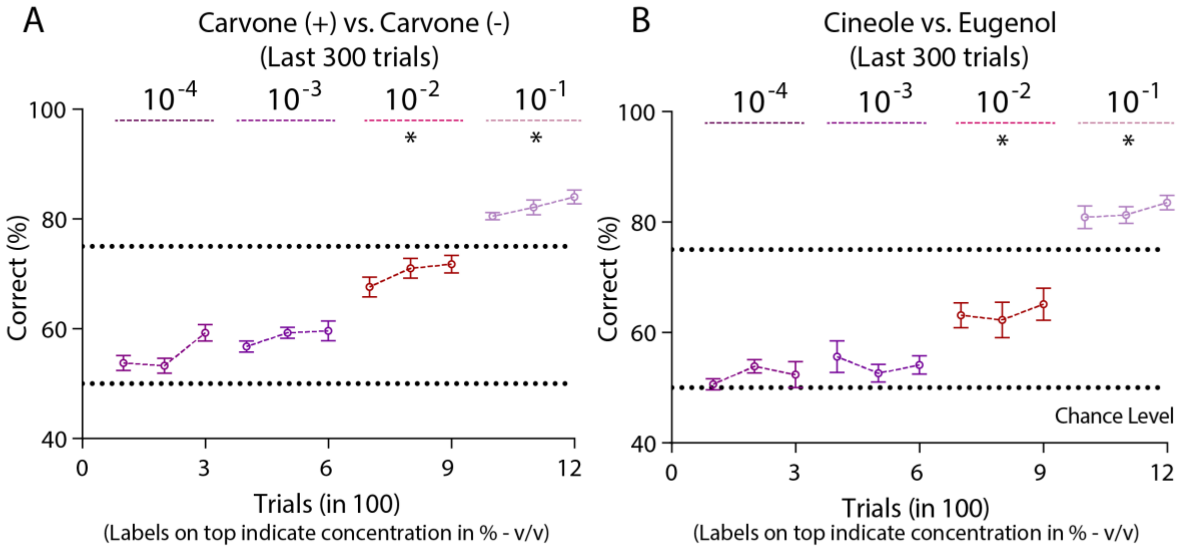


**Figure S7.** Discrimination threshold of animals during odour-based discrimination task

**A.** Animals were trained to discriminate different concentrations of Carvones (+) vs. Carvones (–) to quantify the discrimination threshold. Animals were trained on $10^{-4}$ % (v/v), $10^{-3}$ %, $10^{-2}$ %, and $10^{-1}$ % concentrations. The graphs represent the learning curve for last 300 trials when trained on a particular concentration. The performance of animals was subthreshold (<75%) for all the concentrations except to that of $10^{-1}$ %.

**B.** Animals were trained to discriminate different concentrations of Cineole vs. Eugenol to quantify the discrimination threshold. Animals were trained on $10^{-4}$ % (v/v), $10^{-3}$ %, $10^{-2}$ %, and $10^{-1}$ % concentrations. The graphs represent the learning curve for last 300 trials when trained on a particular concentration. The performance of animals was subthreshold (<75%) for all the concentrations except to that of $10^{-1}$ %.

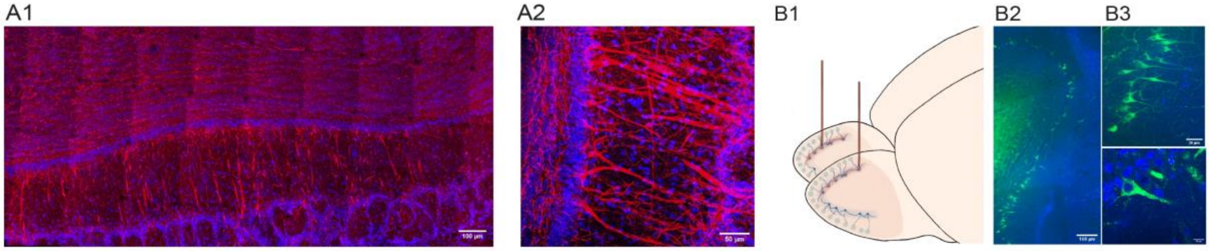


**Figure S8. Piezo2 channels in the olfactory bulb projection neurons**

**A.** **A1.** Tile scan representing the expression of Piezo2 in the coronal section of the mouse olfactory bulb. **A2.** Enlarged image representing the expression of Piezo2 in Mitral cells of the olfactory bulb. Red represents staining corresponding to Piezo2 and blue represents DAPI.

**B.** **B1.** Schematic representing the stereotaxic injections of FM1-43 in the Mitral cell layer of the mouse olfactory bulb. **B2.** Uptake of the FM1-43 dye in the mouse olfactory bulb. **B3.** Enlarged images representing the uptake of FM1-43 (dye specific for Piezo2) in the Mitral cells in the olfactory bulb. Green represents fluorescence from FM1-43 whereas blue represents DAPI.
